# CCR1 inhibition sensitizes multiple myeloma cells to glucocorticoid therapy

**DOI:** 10.1101/2024.12.05.626954

**Authors:** Bert Luyckx, Maaike Van Trimpont, Fien Declerck, Eleni Staessens, Annick Verhee, Sara T’Sas, Sven Eyckerman, Fritz Offner, Pieter Van Vlierberghe, Steven Goossens, Dorien Clarisse, Karolien De Bosscher

**Author notes:** Corresponding author: Correspondence to Karolien De Bosscher. These authors contributed equally and jointly supervised this work: Steven Goossens, Dorien Clarisse, Karolien De Bosscher.

## Abstract

Glucocorticoids (GC) are cornerstone drugs in the treatment of multiple myeloma (MM). Because MM cells exploit the bone marrow microenvironment to obtain growth and survival signals, resistance to glucocorticoid-induced apoptosis emerges, yet the underlying mechanisms remain poorly characterized. Here, we identify that the chemokine receptor CCR1, together with its main ligand CCL3, plays a pivotal role in reducing the glucocorticoid sensitivity of MM cells. We show that blocking CCR1 signaling with the antagonist BX471 enhances the anti-MM effects of the glucocorticoid dexamethasone in MM cell lines, primary patient material and a myeloma xenograft mouse model. Mechanistically, the drug combination shifts the balance between pro- and antiapoptotic proteins towards apoptosis and deregulates lysosomal proteins. Our findings suggest that CCR1 may play a role in glucocorticoid resistance, as the GC-induced downregulation of CCR1 mRNA and protein is blunted in a GC-resistance onset model. Moreover, we demonstrate that inhibiting CCR1 partially reverses this resistance, providing a promising strategy for resensitizing MM cells to GC treatment.

## Introduction

Multiple Myeloma (MM) is a hematological malignancy characterized by clonal expansion of terminally differentiated plasma cells in the bone marrow. The current standard of care for newly diagnosed patients are combinations of immunomodulatory agents (IMIDs, e.g. lenalidomide), proteasome inhibitors (PIs, e.g. bortezomib), monoclonal antibodies (e.g. daratumumab) and glucocorticoids (GCs, e.g. dexamethasone). With the advent of bispecific antibodies (e.g. teclistamab), BiTE (bispecific T-cell engager, e.g. teclistamab-cqyv) and CAR-T based therapies (e.g. ciltacabtagene autoleucel) in the relapsed setting, outcomes for MM patients are gradually improving^1–5^. Despite these treatment advances, MM remains largely incurable due to patients relapsing with resistant disease^6^. Therapy resistance, often in combination with quality-of-life compromising side effects, especially surfaces with prolonged, high-dose GC treatment. Dexamethasone (Dex) is nonetheless a critical treatment pillar and is present in nearly all treatment stages^1^. Dex activates the glucocorticoid receptor (GR) to trigger direct apoptotic effects on malignant lymphoid cells^2–4^. Clinically, Dex augments the response rates of other anti-MM drugs and mitigates side effects of the latter^5^. Altogether, these arguments underscore the ongoing need for further mechanistic understanding that contributes to advances in GC treatment of MM.

Although the bone marrow (BM) microenvironment plays a key role in myeloma pathology, its role in the emergence of GC resistance remains understudied. MM cells exploit the BM niche, where bone marrow stromal cells provide survival and growth signals, either via direct cell-adhesion or via soluble factors^7–9^. For example, SDF-1α^10^ and IL-6^11^ were shown to induce resistance to Dex-induced apoptosis in MM cells. Another soluble factor, CCL3 (macrophage inflaammatory protein-1 alpha, MIP-1α), is one of the main chemokines activating CCR1 (C-C motif chemokine receptor 1)^12^. CCR1 is a GPCR (G Protein-Coupled Receptor) that is highly expressed on the plasma membrane of MM cells and cells within the bone marrow niche, like osteoclasts, osteoblasts, and bone marrow stromal cells^13,14^. As CCL3 itself is produced by bone marrow stromal cells, but also by MM cells themselves, its proliferative effects are conferred in a paracrine and autocrine fashion^15^. Serum levels of CCL3 were shown to correlate with myeloma bone disease and survival^16^, while the CCR1 autocrine loop contributes to decreased sensitivity to the alkylating agent melphalan and bortezomib^17,18^. Preclinical research indicated that blocking CCR1 signaling with antagonists can reduce tumor burden, MM cell dissemination, and osteolytic lesions^13,19,20^. The therapeutic potential of targeting CCR1 in myeloma is therefore high and was recently reviewed^21^.

The current evidence linking the CCR1-CCL3 axis to GCs in MM comes from microarray data in MM cells, showing downregulation of cytokines such as CCL3 by Dex, likely due to the inhibition of NF-κB- mediated expression of CCL3^22^. Adding the fact that blocking CCR1 signaling enhances the sensitivity to other anti-myeloma drugs^17,18^ and taking into account that clinically, GCs augment the response rates to other anti-myeloma drugs^5,23–25^, we hypothesized that targeting the CCR1-CCL3 axis in combination with Dex could potentiate GR-mediated anti-myeloma activity.

The present study investigates the anti-myeloma effects of targeting CCR1 combined with GCs in MM cell lines, primary patient material, and a myeloma xenograft mouse model. We assess the role of CCR1 expression in development of GC resistance and elucidate the mechanism of GC-CCR1 antagonist drug interaction in an unbiased way.

### Materials and methods Cell culture and reagents

Myeloma cell lines were cultured in RPMI1640 GlutaMax medium supplemented with 10% Fetal Bovine Serum, 100 U/mL penicillin and 0.1 mg/mL streptomycin. MM1.S, MM1.R were purchased from ATCC, OPM-2 cells from DSMZ, L363 cells were provided by Prof. M. Engelhardt (Uniklinik Freiburg, Germany). Cells were regularly tested for mycoplasma infection status (LT07-518, MycoAlert kit, Lonza). All experiments were performed in charcoal-stripped serum (CTS, 10706143, Gibco). For *in cellulo* experiments, dexamethasone (D4902, Sigma) and BX471 (HY-12080, Medchemexpress) were dissolved in EtOH as solvent, chloroquine (CQ, C-6628, Sigma) was dissolved in H2O. For *in vivo* experiments, Dex was formulated as dexamethasone-sodiumphosphate (Alfadexx, Alfasan) and diluted in PBS, BX471 was dissolved in propylene glycol. Recombinant CCL3 was ordered from Peprotech (MIP-1α, 300-08).

### Cell viability assays

Cells were seeded in triplicate and treated for the indicated times, after which cellular viability was measured by luminescence-based readout by the CellTiterGlo assay (G7571, Promega). Solvent-treated cells served as controls, luminescence of treated wells was compared to the luminescence level of control wells and expressed as % viability. All graphs depict the average viability of 3 independent biological replicates + SD unless otherwise stated.

### Patient-derived MM cells

Sample acquisition was approved by the ethical commission of the Ghent University Hospital (EC UZG 2018/0906), and informed consent was obtained from all patients. Bone marrow aspirates were filtered through a cell strainer and mixed with a RosetteSep human MM cell enrichment cocktail (15129, Stemcell Technologies). Afterwards, bone marrow aspirates were diluted 1:1 with PBS (+ 2% FBS) and layered on a Lymphoprep gradient using SepMate tubes (Stemcell Technologies). After centrifugation, the cells were washed twice with PBS (+ 2%FBS) and with a red blood cell lysis buffer (0.8% NH4Cl, 0.1 mM EDTA, Stemcell technologies). Thereafter, the enriched MM cells were resuspended in RPMI1640 GlutaMAX (+ 10% CTS) and subjected to a CellTiterGlo cell viability assay.

### Western blots

MM cell lines were treated for the indicated times, washed 2 times with PBS and pelleted. Whole cell lysates were prepared by adding Totex protein lysis buffer (for composition see^26^), denatured, loaded on SDS-PAGE gels and blotted on nitrocellulose membranes. Supplementary Table 7 lists the primary antibodies. Depending on the target and separation of the gel, GAPDH or Tubulin were used as loading control. Secondary antibodies are HRP-conjugated (NA931, NA934, GE-Healthcare). Blots were imaged using the Amersham 680 (GE healthcare) imaging system.

### RT-qPCR

RNA was extracted from cell pellets using the RNeasy mini kit (74106, Qiagen). Equal amounts of total RNA were used for Reverse Transcription (RT) with the iScript cDNA synthesis kit (1708891, BioRad). qPCR reactions were performed using Lightcycler 480 SYBR Green I Master mix (4887352001, Roche), with primers sequences described in Supplementary Table 7. SDHA, YWHAZ, RPL13A served as reference genes for Cq-value normalization. Cq values were analyzed using qBasePlus (Biogazelle).

### Flow cytometry

As a measure for apoptosis, 200 000 cells were stained with eBioscience Annexin V Apoptosis detection kit (88-8006-74, eBioscience). After gating for single cells (Supplementary Fig. 6), the percentage of Annexin V+ cells from the parent population served as apoptotic cell %. CCR1 expression levels were quantified by flaow cytometry after immunostaining with an anti-human CCR1 antibody (MAB145, R&D systems) and a secondary anti-rabbit AlexaFluor 647 antibody. Downregulation of CCR1 after dexamethasone treatment was assessed by relative CCR1 mean flauorescence intensity (MFI) compared to MFI of solvent treated cells. For mice samples, a red blood cell lysis step was performed before CD138-PE staining (60003PE, Stemcell technologies) to identify human myeloma cells.

### Knockdown and overexpression experiments

CCR1 knockdown was done by lentiviral transduction with pLKO.1 vectors containing a shRNA sequence against the CCR1 CDS (TRCN0000008186, TRCN0000008188, BCCM/GeneCorner, http://genecorner.ugent.be) versus a control scrambled shRNA sequence (LMBP 7272, BCCM/GeneCorner). shRNA sequences can be found in Supplementary Table 8. An EGFP-CDS was cloned into the pLKO.1 vector, to allow FACS-sorting of shRNA expressing cells on EGFP. Cell viability assays were performed on sorted EGFP+ populations, 3 days after transduction. Overexpression of CCR1 was done by lentiviral transduction with an in-house constructed lentiviral expression vector (CCR1-OE), using the Golden Gateway cloning method developed by the lab of Prof. Dr. Sven Eyckerman^27^. Brieflay, the plasmid contained the CCR1 CDS (NM_001295.3) coupled to T2A-EGFP, all under control of an EF-1a promoter. T2A is a self-cleaving peptide, which causes ribosomal skipping upon translation. Consequently, CCR1 and EGFP are translated at equimolar amounts, but are physically separated in the cell. MM1.S cells were lentivirally transduced with the either the CCR1-OE plasmid or a control mCherry-OE plasmid. 3 days after transduction, the transduced cells were sorted for EGFP expression. On this sorted population, cell viability assays were performed to assess GC sensitivity.

### *In cellulo* GC-resistance onset model

MM1.S cells were treated for a total of 4-weeks with 0.01 µM Dex, by subculturing and refreshing Dex- medium once every week. Prior to treatment, cells were cultured for one week in RPMI1640 GlutaMAX + 10% CTS to avoid any confounding effects of other steroids and hormones present in the culture medium which contains FBS. GC-sensitivity was assessed by staining for Annexin V and flaow cytometry after treatment with 1 µM Dex for 72h. Gene expression in response to GCs was evaluated by treating cells for 6h with a Dex-range (0.01-1 µM) and subsequent RNA extraction and qPCR for target genes. Regarding GC-responses on protein levels, the cells were treated with a Dex-shot (1 µM) for 24h, after which a staining for CCR1 on the plasma membrane was done as described above. CCR1 expression level was expressed as % MFI from solvent treated control at the respective timepoint when the assay was performed.

### Myeloma xenograft mouse model

NXG (005557, The Jackson Laboratory) female mice (9-weeks age, n=24) were injected i.v. with 5.10^6^ MM1.S-luciferase expressing cells in 100 µL PBS. Engraftment was confirmed by measuring bioluminescence using the IVIS Lumina II imaging system (PerkinElmer). 5-weeks after engraftment, mice were randomized on bioluminescence level into vehicle (propylene glycol (PG) + PBS, n=6), Dex (n=6), BX471 (n=6) and Dex+BX (n=6) treatment groups. Dex was administered i.p. daily at 2.5 mg/kg in PBS and BX471 was i.p. injected every 12 hours at 50 mg/kg in propylene glycol. All mice received an equal amount of propylene glycol and PBS to correct for solvent-induced effects. Mice were treated for 48 hours, further treatment was halted due to general health concerns of the mice, attributed to solvent-related toxicity (see Supplementary Table 3 for a detailed description). During the experiment, total myeloma burden was measured via bioluminescence. Only living mice, or mice producing a reliable BLI signal (average radiance > 3×10^4^ p/s/cm²/sr) were included in the analysis (Solvent n=6, BX n=3, Dex n=6, Dex-BX n=5). Upon sacrifice, bone marrow and blood were collected and myeloma burden was assessed by flaow cytometry after CD138-PE staining (60003PE, Stem Cell technologies). Mice blood parameters were measured with a Vetscan HM5 Hematology Analyzer (Abaxis). Serum mouse OPG levels were determined by ELISA (RAB0493, Sigma). All *in vivo* experiments were approved by the ethical committee on animal welfare at Ghent University Hospital (ECD 23-22).

## Results

### Increased CCR1 expression during GC-based treatment is associated with poor prognosis

We first investigated whether CCR1 expression had prognostic value in terms of overall-survival (OS) in GC-treated myeloma patients. To this end, we used public RNA-sequencing data of bone marrow samples of myeloma patients included in the CoMMpass-trial, 740/750 (98.7%) of which received GC-based treatment (Dex or Prednisolone) as first-line therapy (Supplementary Table 1). We found that CCR1 expression was not associated with OS at diagnosis (Fig. 1a, *p* = 0.8). However, as disease progressed, the expression of CCR1 concurrently also increased (Fig. 1b, *p* = 0.045, n = 48 patients with follow-up samples, 47/48 received GC-based treatment (97.9%)). We next subdivided these patients based on the evolution of CCR1 expression. Notably, patients having stable CCR1 expression (CCR1TPM,progression / CCR1TPM,diagnosis < 2) had a markedly better OS than patients with elevated CCR1 expression (CCR1TPM,progression / CCR1TPM,diagnosis ≥ 2, Fig. 1c, *p =* 0.0072).

**Figure 1:**
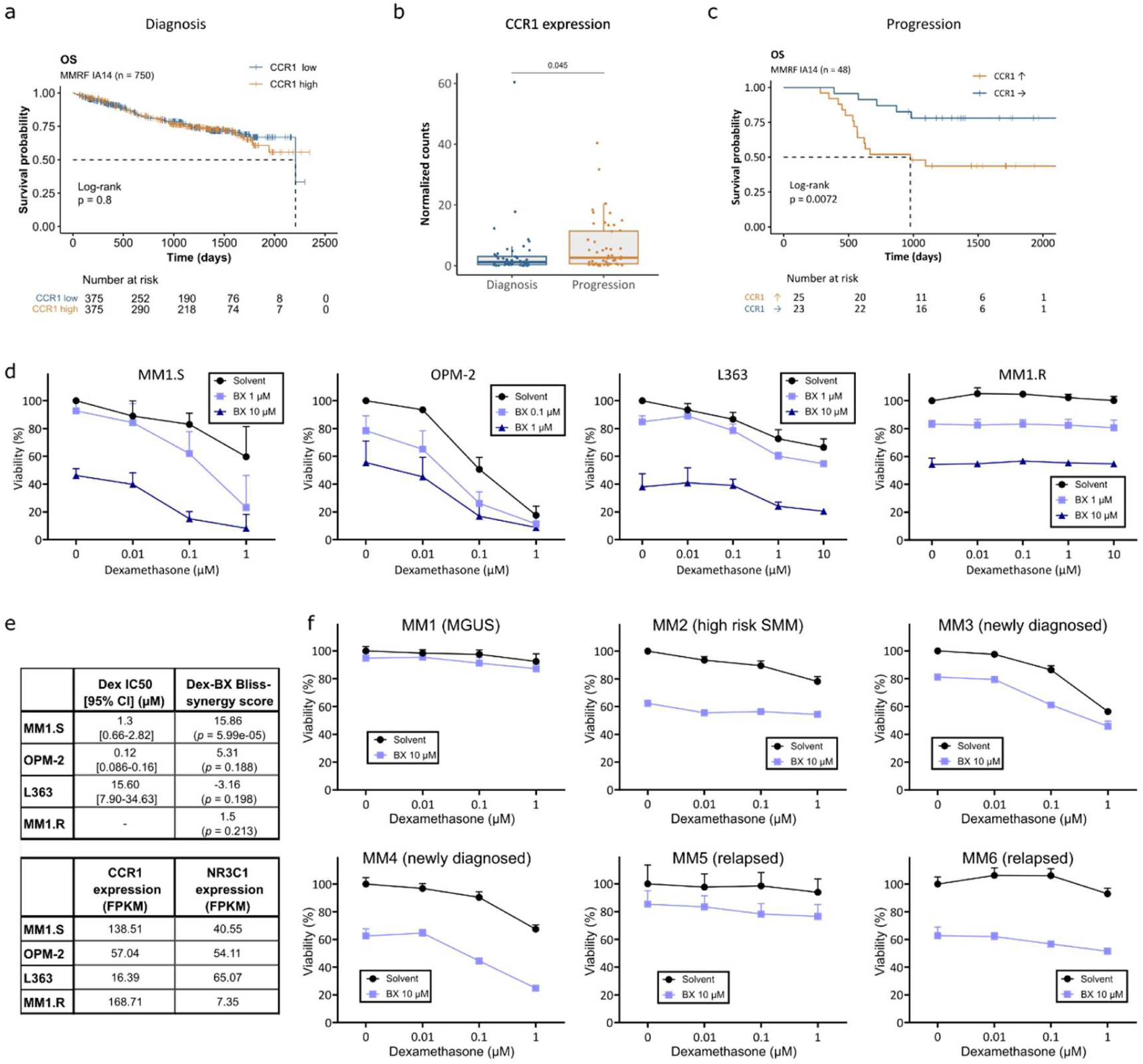
Increased CCR1 expression after a GC-based treatment cycle confers poor prognosis, while inhibition of CCR1 signaling with BX471 enhances GC sensitivity of myeloma cells. a OS of patients included in the MMRF CoMMpass trial (IA14, n=750), stratified on CCR1 expression of bone marrow samples upon diagnosis. b CCR1 transcript expression levels of MM patients with follow- up RNA sequencing in the CoMMpass dataset (IA14, n=48). c OS of MM patients with follow-up RNA sequencing, categorized on stable CCR1 expression (CCR1 →, CCR1TPM,progression / CCR1TPM,diagnosis < 2) or increasing CCR1 (CCR1 ↑, CCR1TPM,progression / CCR1TPM,diagnosis ≥ 2) expression. For a,c statistical significance was tested by a log-rank test with a null hypothesis of no difference in survival between both groups. For b, the p-value is shown for a paired t-test. d Viability of myeloma cell lines after 24h treatment with a Dex-range (+BX471). Viability expressed as % versus control, readout by CellTiterGlo. Data are presented as mean viability + SD (n=3). e Dex IC50 values for MM cell lines used in d and Bliss synergy scores for synergy between Dex and BX471. The score represents the average excess response due to drug interactions. A score >10 indicates the interaction between two drugs is likely synergistic, (https://synergyfinder.fimm<u>.fi</u>). IC50 values and synergy scores were calculated based on data shown in d. Reported p-values are for a t-test under the null hypothesis of drug independence. CCR1 and GR (NR3C1) FPKM levels from myeloma cell lines were obtained from Keats lab (https://keatslab.org) f Viability of primary myeloma cells isolated from bone marrow aspirates of MM patients after 48h (MM2) or 72h treatment with a Dex-range (+BX471 10 µM), readout by CellTiterGlo. Data are presented as mean viability + SD from 3 technical replicates. Note that the limited culturing time of primary myeloma cells does not allow for performing biological replicates.

### Inhibition of CCR1 signaling enhances GC sensitivity in MM cell lines and primary myeloma cells

As increasing CCR1 levels during GC-based therapy are associated with poor survival, we examined the effect of inhibiting CCR1 signaling with BX471 (a non-peptide inhibitor, highly specific for CCR1^28^) together with Dex treatment on the myeloma cell viability. Several myeloma cell lines with varying degrees of GC responsiveness (Fig. 1d-e, black curves, ref.^26^) and GR and CCR1 expression levels (Fig. 1e) were used. Adding BX471 to a Dex-concentration range increased the cytotoxicity of Dex (Fig. 1d, blue curves). This combination effect was either synergistic (MM1.S), additive (OPM-2, L363) or absent (MM1.R, lacks GR protein^26^), and thus most pronounced in the GC-sensitive cell lines with high CCR1 mRNA expression levels (Fig. 1e, Supplementary Fig. 1a). BX471 alone only had anti-MM effects at higher concentrations (10 µM) in all cell lines. We also observed GC-mediated CCR1 downregulation, which is likely transcriptionally modulated by GR, given the timing of downregulation in GR expressing cell lines and the absence of CCR1 downregulation in MM1.R (Supplementary Fig. 1b).

To validate the drug combination in patient material, we isolated primary MM cells from bone marrow aspirates of patients at different stages of the disease: MGUS (Monoclonal Gammopathy of Unknown Significance, an MM precursor stage), SMM (Smoldering Multiple Myeloma, a subsequent MM precursor stage), diagnosis or relapse (Fig. 1f, Supplementary Table 2). The Dex-BX combination was most effective in primary cells from newly diagnosed patients, which also still responded to Dex *ex vivo* (MM3 and MM4). BX alone (10μM) also induced killing of cells from different disease stages, although the response varied across patients (5-40% cell killing, Fig. 1f).

### Pharmacological inhibition or knockdown of CCR1 enhances the GC sensitivity of MM cells

To complement the cell viability assays (Fig. 1d) with evidence of direct apoptotic cell killing, we performed Western analysis of apoptotic proteins and Annexin V/7-AAD flaow cytometry experiments (Fig. 2 and Supplementary Fig. 2). We specifically tested whether the drug combination could induce apoptosis. Neither BX471 nor dexamethasone alone induced apoptosis in MM1.S, the model in which we observed most effects (Fig. 1d-e). Remarkably, the Dex-BX471 combination rendered the bulk of the cells (75 %) positive for Annexin V, a hallmark for early-stage apoptosis (Fig. 2a-b). This was further confirmed by cleavage of apoptotic markers caspase 3 and PARP (Fig. 2c). In OPM-2 cells, in which the IC50-value is lower for Dex (Fig. 1e), Annexin V^+^ cells and apoptotic markers were elevated with Dex mono treatment, adding BX471 did not further increase these levels (Supplementary Fig. 2a-c).

**Figure 2:**
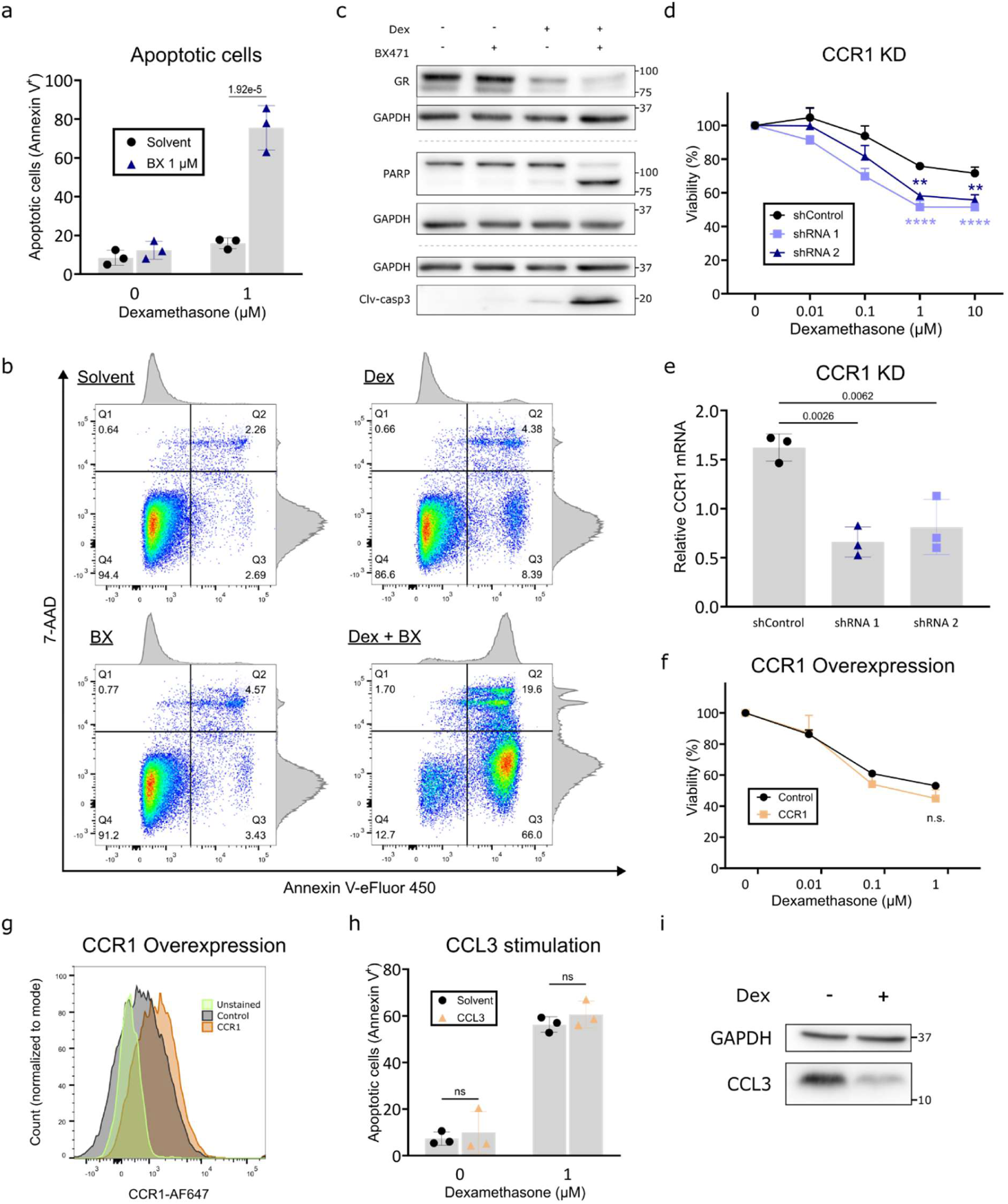
Inhibition of CCR1 signaling with BX471 or knockdown of CCR1 enhances Dex-induced apoptosis of MM1.S cells. a Apoptotic MM1.S cells (Annexin V positive cells) determined by flaow cytometry after 24h treatment with 1 µM Dex (+ BX471 1µM). Reported p-value is for 2-way ANOVA with post-hoc testing (n=3). b Representative Annexin V/7-AAD plot after 24h Dex 1 µM (+ BX471 1 µM) treatment. c Western blot of apoptotic markers of MM1.S whole cell lysates treated for 24h with 1 µM Dex (+ BX471 1 µM), GAPDH served as loading control (n=3) d Viability after 24h treatment with a Dex concentration range. Performed on MM1.S cell populations, sorted for expression of a shRNA-sequence directed against CCR1 (shRNA 1 and 2) versus a scrambled shRNA sequence (shControl). Reported values are mean viability + SD, compared to the respective control (solvent treated) from n=3 independently generated MM1.S CCR1 KD populations. 2-way ANOVA with post-hoc testing was performed. Indicated significance level is for comparisons between shControl and shRNA at a certain Dex-concentration (** corresponds to *p*<0.01 and **** to *p*<0.0001). e CCR1 mRNA levels of the cell populations used in d, reported *p*-value is for 1-way ANOVA with post-hoc testing (n=3). f Viability after 48h treatment with a Dex-range, performed on MM1.S cell populations with CCR1 overexpression versus control. Reported values are mean viability + SD, compared to the respective control (solvent treated) from n=3 independent biological replicates on a MM1.S cell line sorted for expression of a CCR1 CDS construct versus an MM1.S cell line with sorted for overexpression of a control CDS (mCherry). 2-way ANOVA with post-hoc testing was performed for all Dex-concentrations, with the *p*-value indicated for comparison between Control and CCR1 at a Dex = 1 µM. g CCR1 surface protein expression of the populations used in f, as measured by flaow cytometry after staining with a CCR1 primary- and an Alexa- Fluor647 labeled secondary antibody. h Apoptotic MM1.S cells determined by flaow cytometry after 72h treatment with Dex (1µM), in medium supplemented with 50 ng/mL recombinant CCL3 protein. Reported *p*-value is for 2-way ANOVA with post-hoc testing (n=3). i Western blot for CCL3 of MM1.S whole cell lysates, treated for 24h with 1 µM Dex. GAPDH as loading control (n=3).

To further elucidate the impact of CCR1 expression on GC sensitivity of MM1.S cells, we performed shRNA-mediated knockdown of CCR1. In these cells, lower expression levels of CCR1 (± 45% reduction) led to a significant increase in GC sensitivity (Fig. 2d-e). Conversely, overexpression of CCR1 did not significantly alter Dex-sensitivity of MM1.S cells (Fig. 2f-g), neither did stimulation of CCR1 signaling by exogenous CCL3 (Fig. 2h). However, this might be due to saturation, given the already substantial endogenous CCL3 production in MM1.S cells (Fig. 2i). In contrast, in OPM-2 cells, which have lower levels of endogenous CCL3, there was a trend to reduced sensitivity to Dex-induced apoptosis, when CCL3 was added to the medium (Supplementary Fig. 2d-e).

### Dex and BX471 combination lowers tumor burden in a mouse myeloma xenograft model and has positive impact on myeloma bone disease markers

Next, we evaluated whether the combination treatment would show anti-myeloma effects *in vivo*. Hereto, we generated MM1.S cells stably expressing a transgene EGFP-luciferase reporter construct, which were subsequently i.v. injected in NXG immunodeficient female mice (Fig. 3a). After evidence of myeloma engraftment and progression (Supplementary Fig. 3a), mice were randomized in 4 groups (n=6/group) based on bioluminescent imaging (BLI) and treated with either Dex (2.5 mg/kg daily), BX471 (50 mg/kg twice daily), Dex-BX471 combination or vehicle (solvent control). After two days of treatment, a significant increase in BLI signal was observed in all vehicle-treated control animals, indicative of progressive disease. While no effects of BX471 alone was seen on in vivo myeloma growth, the combination therapy increased the Dex therapeutic effects as evidenced by total BLI, in combination with a significant reduction in the levels of MM cells in the bone marrow (Fig. 3c, *p =* 0.0182), and a trend to decreased circulating MM cells compared to any other treatment groups (Fig 3d, n.s.). As a measure for myeloma bone disease, the osteoprotegerin (OPG) levels in blood serum were determined, which are inversely correlated with the degree of myeloma bone disease^29^. OPG levels were significantly elevated by the combination treatment (*p*=0.0084 and *p*=0.0168 compared to solvent- and BX-treated mice, respectively), supporting a reduction in myeloma bone disease. The combination therapy itself was well tolerated with no differences in mouse weight and vital blood parameters (Supplementary Fig. 3b-c).

**Figure 3:**
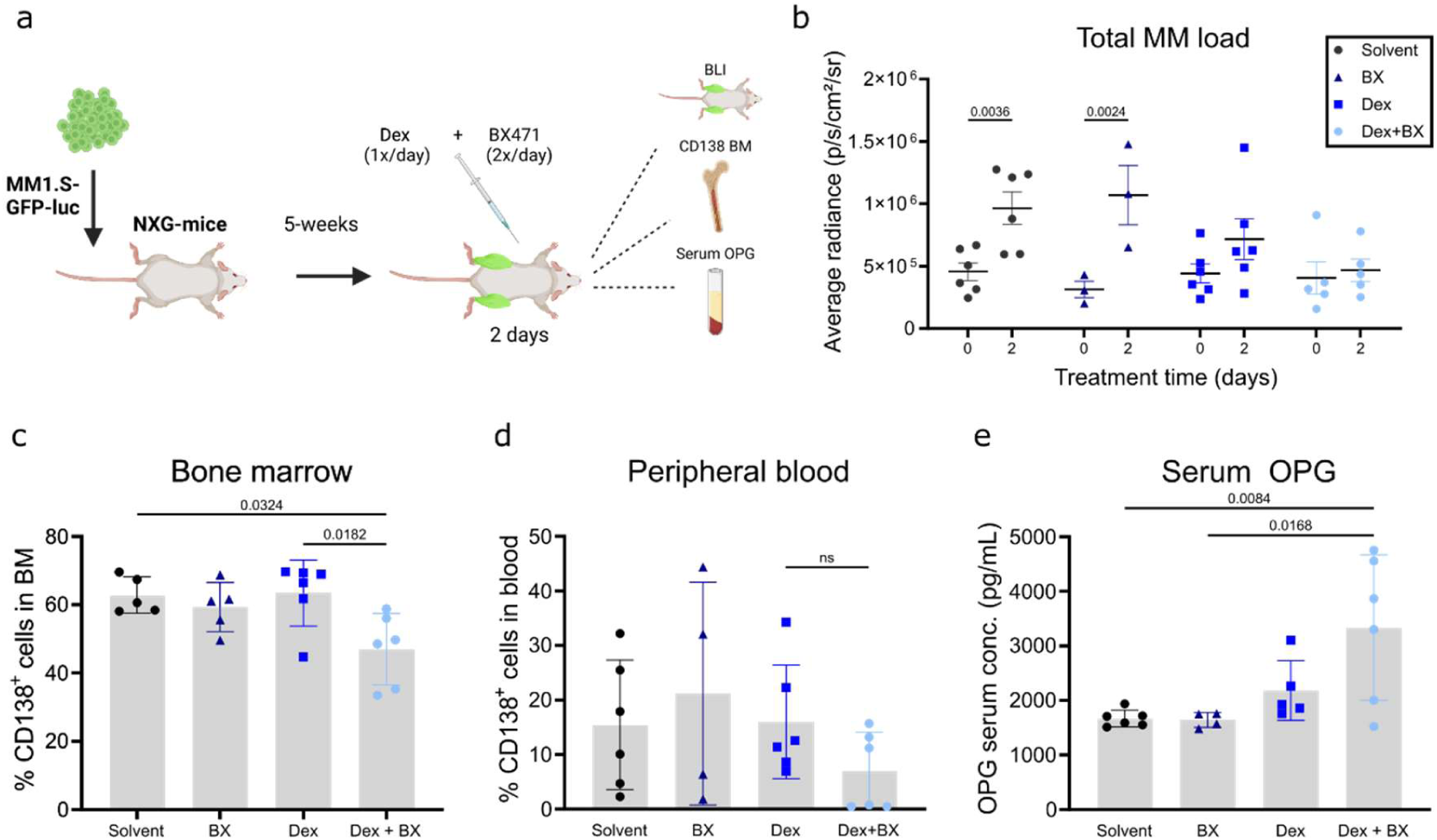
Dex and BX471 combination therapy leads to lower levels of MM cells in the bone marrow and higher levels of serum OPG in mice. a Overview of the MM1.S-xenograft NXG mice model. A luciferase expressing MM1.S cell line is injected intravenously in female NXG mice (n=6 per group) and after tumor establishment, mice are randomized on total BLI signal and treated 1x/day with 2.5 mg/kg Dex and/or 2x/day with 50 mg/kg BX471 for 2 days in total. b Quantification of total MM tumor load at start (day 0) and end of therapy (day 2). Mice in which no BLI signal was generated (< 3×10^4^ p/s/cm²/sr average radiance) or died during anesthesia were excluded (solvent n=6, BX n=3, Dex n=6, Dex + BX n=5). c, d Percentage of CD138^+^ myeloma cells in the bone marrow and peripheral blood of NXG mice at the end of therapy. Analyzed by flaow cytometry after staining with an anti-CD138-PE antibody. e Serum mouse OPG levels at the end of therapy, measured by ELISA. Two BX-treated mice were excluded from the analysis, as no blood was collected during sacrifice (solvent n=6, BX n=4, Dex n=6, Dex + BX n=6). For b 2-way ANOVA and for c-e 1-way ANOVA with post-hoc testing was performed. *p-*values for statistically significant differences are shown (*p*<0.05), non-significant differences are indicated by ns or omitted.

### GC resistance is associated with loss of GC-induced CCR1 downregulation, while BX471 can resensitize partial GC-resistant MM cells to Dex

As CCR1 expression levels in MM patients increase under GC-based treatment regimens as disease progresses (Fig. 1b), we wondered whether CCR1 expression could be associated with a GC-resistant phenotype. We constructed an *in cellulo* GC resistance model in MM1.S cells that mimics the onset of GC resistance by culturing MM1.S cells for 4-weeks with low dose (0.01μM) Dex, or solvent control (Fig. 4a). After 1-week of Dex treatment, cells are still sensitive to Dex-induced apoptosis, however after 4-weeks, MM1.S cells become partially resistant to GC-induced apoptosis (Fig. 4b, *p*=2.5e-6). We found that this partial GC resistance, established at 4 weeks of low-dose Dex treatment, could be reversed by combining Dex with BX471, hereby effectively sensitizing MM1.S cells to GCs (Fig. 4c-d). Notably, the Dex response of these partial GC-resistant cells that were additionally treated with BX is no longer different from the Dex response of the GC-sensitive control population that also received BX471 (Fig. 4d).

**Figure 4:**
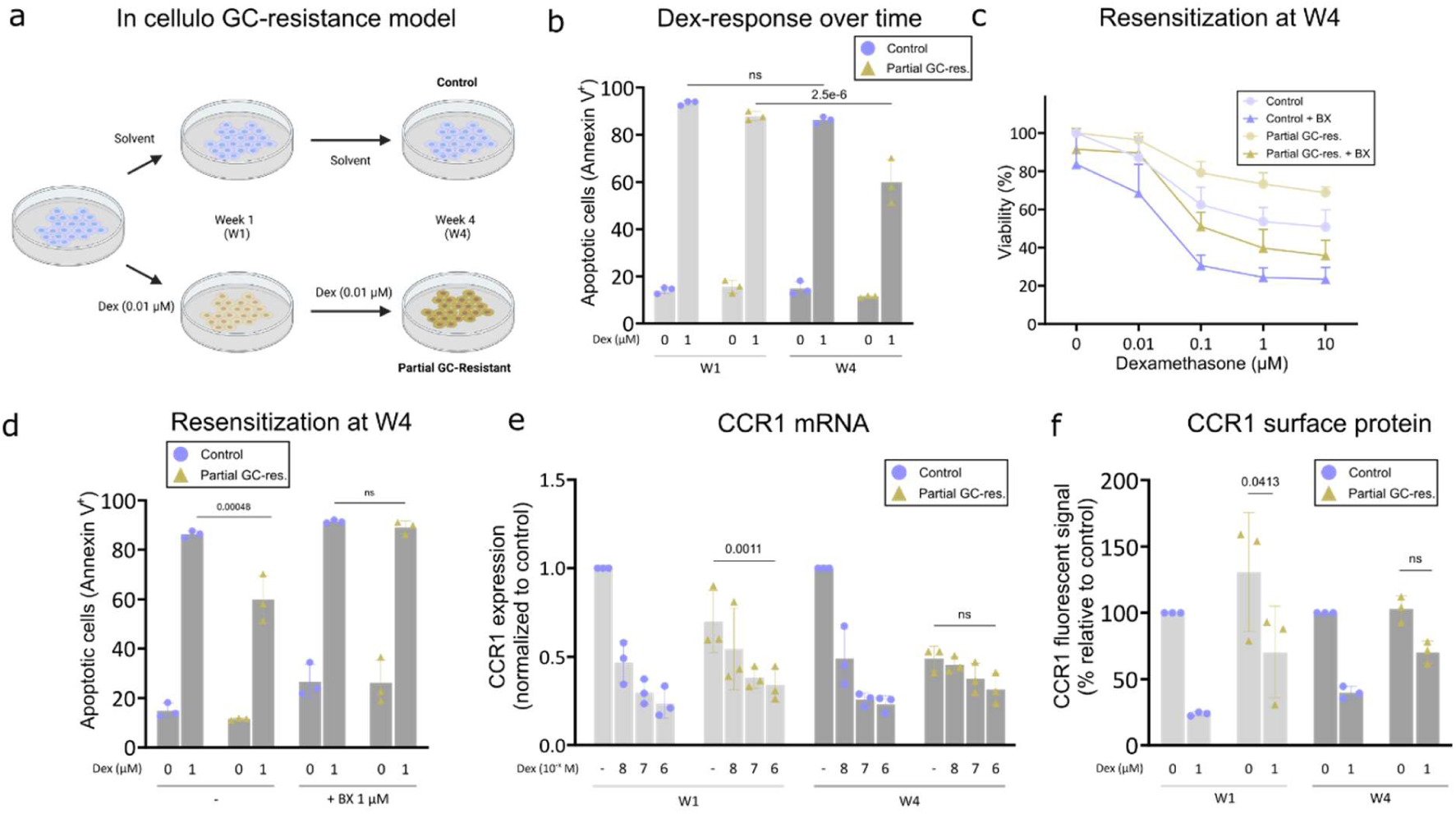
Partial GC-resistance can be reversed by BX471 and is associated with loss of GC-induced CCR1 downregulation. **a** Schematic overview of the *in cellulo* GC-resistance model. MM1.S cells are treated for 4-weeks with low-dose Dex (0.01μM) to induce partial GC-resistance. **b** Percentage apoptotic MM1.S cells after an additional 72h treatment with 1 µM Dex of the respective control and partial GC-resistant cell populations at W1 and W4 of the model. Apoptotic cells were measured by flaow cytometry after Annexin/7-AAD staining (n=3 biological replicates). **c** Viability of partial GC-resistant versus control MM1.S cells at W4 of the *in cellulo* GC-resistance model after 24h additional treatment with a dexamethasone range (+ BX471 1 µM). Readout via CellTiterGlo (n = 3 biological replicates). **d** Apoptotic cells after 72h treatment with 1 µM Dex (+ BX471 1 µM). Assay was performed on partial GC-resistant cells versus control at W4 of the GC-resistance model. Apoptotic cells were measured by flaow cytometry after Annexin/7-AAD staining (n=3 biological replicates). **e** CCR1 mRNA levels in response to a 6h treatment with a Dex concentration range (0.01 – 1 µM), determined via qPCR. The reported value is the average relative CCR1 expression value of 3 biological replicates of the model (n=3). Values are relative to the CCR1 expression value of the respective control of that biological replicate at the time point of sample collection (= control cells at W1 or W4 respectively, treated 6h with solvent control). **f** CCR1 protein levels in response to a 24h treatment with 1 µM Dex, measured via flaow cytometry after staining for cell surface CCR1 expression with a primary anti-CCR1 antibody and an AlexaFluor-647 labeled secondary antibody. Values are relative MFI to the respective control at the time of staining (W1 or W4 control cells treated 24h with solvent control). Indicated p-values in **b,d,e** and **f** are for 2-way ANOVA with post-hoc testing, n=3 biological replicates.

We subsequently examined whether CCR1 expression changes upon prolonged GC treatment in this GC resistance model. Comparable as observed with parental MM1.S cells (Supplementary Fig. 1b), we found a Dex-induced downregulation of CCR1 mRNA levels (Fig. 4e), which remained relatively consistent over the course of the treatment. However, as GC resistance emerges, the capacity of Dex to downregulate CCR1 is dampened (Fig. 4e, *p*=0.0011 and n.s. at W1 and W4 respectively). These results largely align with our findings on the CCR1 surface protein level, where this dampening effect is similar (Fig. 4f, Supplementary Fig. 4).

### Dex and BX471 combination treatment suppresses lysosome associated proteins and deregulates the pro-and antiapoptotic protein balance in favor of apoptosis

To gain mechanistic insight into how inhibition of CCR1 signaling synergizes with Dex to induce MM cell killing, we performed mass spectrometry-based shotgun proteomics on MM1.S cells treated with Dex (+ BX471). The combination rendered 153 uniquely differentially expressed proteins (*padj* ≤ 0.05, |log2FC| ≥ 1 compared to solvent), as opposed to cells solely treated with Dex (24) or BX471 (1) (Fig. 5a, a full list is available in Supplementary Table 4). The distinct protein expression profiles are also reflaected in the heatmap (Fig. 5b). To fully grasp the differences in protein expression between Dex- and Dex-BX treated cells, we fitted a linear model that incorporated the differential expression of the mono treatments Dex and BX versus that of the Dex-BX combination, a so-called interaction model (Dex:BX). This yielded a total of 109 differentially expressed proteins (*padj* ≤ 0.05, |log2FC| ≥ 1; 82 downregulated and 27 upregulated, Fig. 5c, a full list is available in Supplementary Table 5). Gene set enrichment analysis (GSEA, KEGG 2021 Human pathways) of differentially expressed proteins of the interaction term revealed that genes were significantly enriched for ‘Lysosome’ (*p=*0.0028) and ‘Apoptosis’ (*p=*0.033) (Fig. 5d); the list of proteins contributing to the enriched terms can be found in Supplementary Table 6. Of note, lysosomal proteases from the cathepsin family (CTSA, CTSB, CTSZ) and other lysosomal enzymes (TPP1, NEU1, Supplementary Figure 5a) were significantly downregulated by Dex-BX compared to any other treatment. One of the genes significantly downregulated by the combination treatment and contributing to the ‘apoptosis’ term, is the anti- apoptotic protein Mcl-1.

**Figure 5:**
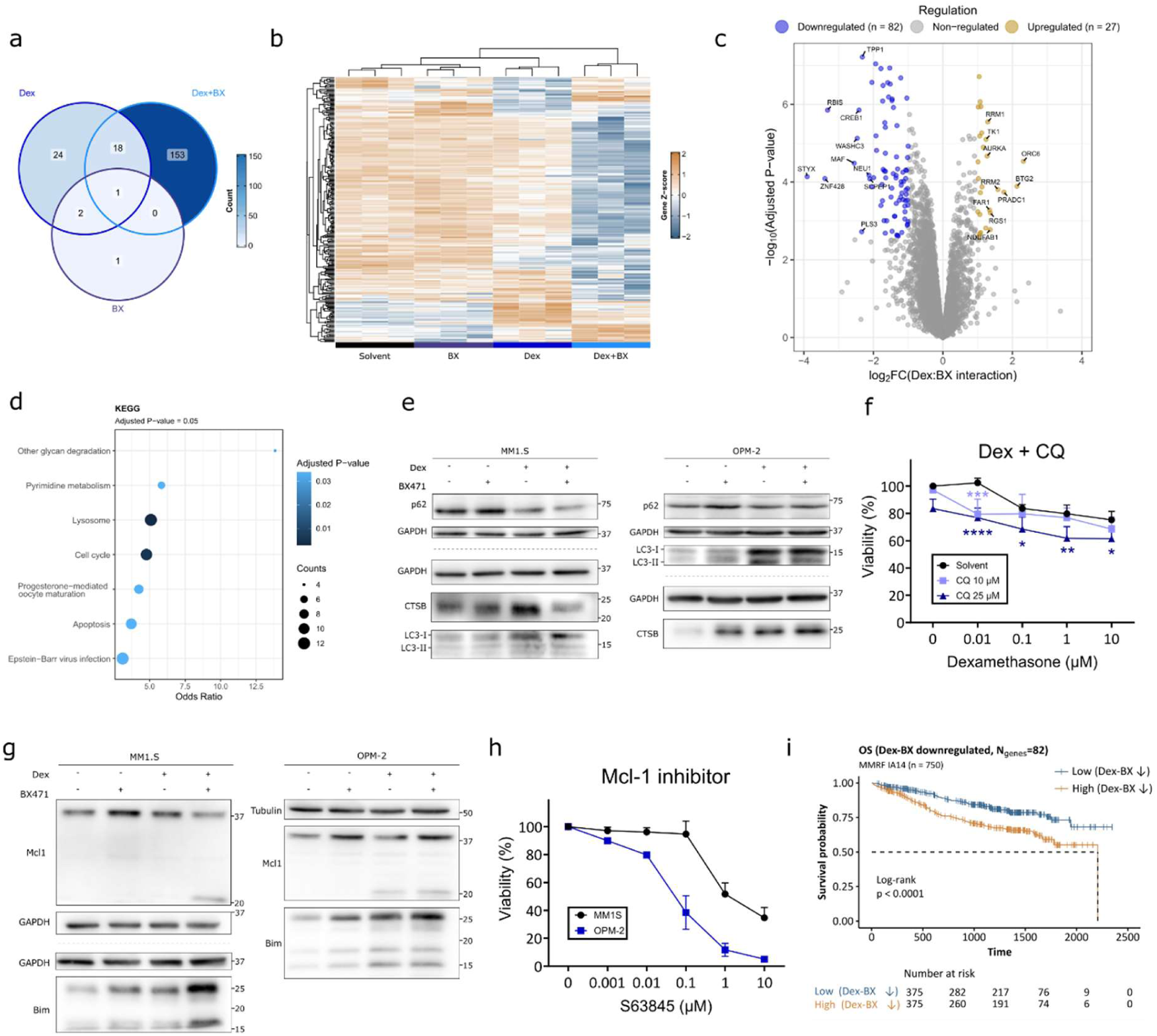
Dex-BX treatment leads to a distinct proteome profile with downregulation of lysosomal proteins and shifted balance of pro- and anti-apoptotic proteins. MM1.S cells were treated for 24h with Dex 1 µM, BX471 1 µM or a combination. Cell pellets were processed to allow for mass spectrometry-based shotgun proteomics. **a** Venn diagram showing the amount of differentially expressed proteins in each treatment group (|log2FC| ≥ 1 and *padj*-value ≤ 0.05) **b** Heatmap and clustering of MaxLFQ intensity Z-scores of all differentially expressed proteins (*padj*-value ≤ 0.05). **c** Volcano plot of differentially expressed proteins of the Dex:BX interaction term, with colored dots indicating proteins with |log2FC|≥ 1 and *padj*-value ≤ 0.05. Differential expression analysis was done by fitting a linear model using the R limma package while considering Dex + BX as a distinct treatment for **a,b**, while for **c** Dex + BX was considered as an interaction Dex:BX between two treatments Dex and BX. **d** GSEA-analysis for KEGG pathway, based on all differentially expressed proteins of the interaction term Dex:BX (*padj*-value ≤ 0.05). **e, g** Western blot of MM1.S and OPM-2 cell lysates treated for 24h with Dex 1 µM (+ BX471 1 µM), GAPDH or Tubulin as loading control (n=3). **f** Viability of MM1.S cells treated for 24h with a Dex-range (+ CQ), readout via CellTiterGlo. Viability is expressed relative to respective control (0 µM Dex) and is the mean + SD of 3 biological replicates. 2- way ANOVA with post-hoc testing was performed. Indicated * denote the significance level of the comparison of CQ10 and CQ25 viability with solvent control at a certain Dex-concentration (* corresponds to *p*<0.05, ** to *p*<0.01, *** to *p*<0.001 and **** to *p*<0.0001). **h** Viability of MM1.S and OPM-2 cells after 24h treatment with a S68345 range, readout via CellTiterGlo. Viability is expressed relative to solvent control and is the mean + SD of 3 biological replicates. **i** OS of patients included in the MMRF CoMMpass trial (IA14, n = 750), stratified on expression of Dex:BX interaction downregulated genes in bone marrow samples of patients at diagnosis. Statistical significance was tested by a log-rank test with a null hypothesis of no difference in survival between both groups.

### Lysosomal protease levels are lowered by Dex-BX combination, while Dex induces autophagy

To further elucidate the role of the lysosome and lysosomal proteases in the action mechanism of the Dex-BX combination treatment and MM cell apoptosis, we examined how lysosomal markers and markers of autophagy respond to Dex-BX in MM1.S and OPM-2 cells. P62 (SQSTM1) is selectively degraded by autophagy, and its total cellular levels inversely correlate with autophagic activity^30^. We found that Dex and the Dex-BX combination treatment reduce p62 protein levels in MM1.S, but not in OPM-2, cells (Fig. 5e). BX471 treatment by itself has little to no impact. LC-3 serves as another proxy for autophagic activity, with accumulation of LC-3 II (the lipidated form of LC-3 I) denoting an accumulation of autophagosomes and impairment of the autophagic process, as LC-3 II gets degraded by autophagy^31^. In MM1.S cells, there is mainly an increase in the LC-3 I form with Dex(-BX) treatment and no accumulation of LC-3 II. Along with the reduction in p62 levels, this indicates increased autophagic activity, largely driven by Dex. In OPM-2 cells, both LC-3 I and II are upregulated by Dex(- BX), again indicating increased autophagic activity, induced by Dex. In contrast, in MM1.S but not in OPM-2 cells, only the combination treatment reduced the protein levels of the lysosomal protease cathepsin B (CTSB, Fig. 5e), in this way leading to a reduction in autophagic clearance capacity of the cells and likely contributing to the observed enhanced cell killing. This reasoning is strengthened by an experiment wherein CQ (chloroquine), an agent blocking lysosomal degradation, was combined with Dex and resulted in an increased cytotoxic effect (Fig. 5f).

### Antiapoptotic Mcl-1 is downregulated by Dex-BX combination, while proapoptotic Bim is upregulated

One of the terms contributing to the KEGG ‘Apoptosis’ term is Mcl-1, an antiapoptotic protein, which is downregulated by Dex-BX combination in MM1.S and by Dex itself in OPM-2 (Fig. 5g). A lower migrating protein detected by the Mcl-1 antibody around 20 kDa likely indicates the degraded form of Mcl-1^32^ (Fig. 5g). That MM cells are highly dependent on Mcl-1 for survival^33,34^, was confirmed by the cytotoxic effects of S63845, a specific Mcl-1 inhibitor (Fig. 5h). Additionally, proapoptotic Bim (BCL2L11) typically gets upregulated by Dex and is necessary for Dex-induced apoptosis of lymphoid cells^2^. In MM1.S and OPM-2 cells, Bim is upregulated by Dex and even more so by Dex-BX. Aside from its effects on lysosomal proteases (Fig. 5e), Dex-BX also shifts the balance between pro- and antiapoptotic proteins (Fig. 5g), collectively explaining the cytotoxic effects of Dex-BX combination therapy.

Finally, to shed light on clinical relevancy of the altered proteome of Dex-BX treated MM cells, we analyzed whether the expression levels of Dex-BX regulated proteins had predictive value in terms of PFS and OS in the CoMMpass cohort of 750 myeloma patients. Because the latter contains transcriptomics data, the gene counterparts of the proteins downregulated by the Dex:BX combination term (Ngenes=82, *padj*≤0.05, |log2FC|≥ 1, Supplementary Table 5) were used as input for survival analysis. Patients with low expression of these genes had significantly better PFS and OS (Fig. 5i, Supplementary Figure 5b). Conversely, when the analysis was done with genes upregulated by the Dex:BX combination term (Ngenes=27), again patients having low expression of these genes, had significantly worse PFS and OS than patients with high expression of this gene set (Supplementary Figure 5c-d).

## Discussion

In this study, we demonstrate the contribution of CCR1 in determining the response to GCs in MM cells. By using the CCR1-specific, competitive antagonist BX471^28^, we show that blocking of CCR1 signaling can enhance the cytotoxicity of GCs in MM cell lines, primary patient samples and *in vivo*. Prior research indicated that stimulation of CCR1 with its main ligand CCL3 affects cell signaling pathways controlling growth, survival and migration^10,15,35^. Accordingly, it is conceivable that inhibition of CCR1 signaling by BX471 enhances the apoptotic effects of Dex (Fig. 2a-c). Furthermore, knockdown of CCR1 itself, hereby reducing constitutive signaling by CCR1 which can adopt an active state even in absence of the ligand^36^, also further sensitized MM1.S cells to GCs (Fig. 2d-e). Further stimulation of CCR1 signaling by exogenous CCL3 or overexpression of CCR1 did not reduce GC-induced apoptosis. Our findings align with previous reports, where CCL3 could not overcome Dex-induced growth inhibition^15^. One potential explanation is that the high endogenous CCL3 expression of MM1.S cells^37^ saturates the system and hereby limits further stimulation. At the same time, the high autocrine levels of CCL3 in MM1.S cells highlight the dependency of this cell model on CCL3. They can be a contributing factor as to why Dex-BX synergism was only observed in MM1.S cells, and not in OPM-2 or L363 cells, where only additive effects were noted. The coincident high expression of both CCR1 and GR (Fig. 1e), the respective targets of BX471 and Dex, could be another explanation for the synergism in MM1.S and its absence in MM1.R. In primary patient samples, differences in GC- and BX471 sensitivity were observed (Fig. 1f). However, no data on CCR1 levels could be generated due to too limited cell numbers. Nevertheless, our *in silico* analyses of the MMRF CoMMpass RNA-sequencing dataset demonstrated that relapsed patients, which are generally GC-resistant, exhibit overall a higher CCR1 expression (Fig. 1b). These findings may indirectly explain our *ex vivo* data, where we found that in cells from relapsed patients, BX471 alone still had anti-myeloma properties. In addition, recent reports pointed out that a CCL3 neutralizing antibody enhances cytotoxicity of melphalan^17^ and that CCR1 expression was associated with decreased sensitivity to bortezomib^18^, further highlighting the relevance of the CCL3-CCR1 axis in a broader MM drug-sensitivity perspective.

In the MMRF CoMMpass RNA-sequencing dataset, CCR1 expression levels upon diagnosis are not correlated with patient prognosis, contrary to what was reported from other micro-array based datasets^35^. However, when CCR1 expression increases under GC-based therapy, then it does become a prognostic factor negatively impacting OS. We have shown using our *in cellulo* GC-resistance model, that partial resistance to GC-induced apoptosis is associated with loss of Dex-induced downregulation of CCR1. This progression of CCR1 expression under pressure of GC-based therapy in patients, may reflaect the loss of GC-induced CCR1 downregulation due to the development of partial resistance to GCs. Hence, this could serve at least as a partial explanation for the worse prognosis of these patients. Since a loss of CCR1 downregulation was thus far only observed in GC-resistant MM1.S cells, future independent analysis of CCR1 protein levels of paired patient samples before and after GC-treatment may help to support this hypothesis further. Additional proof that CCR1 expression is a factor determining GC-resistance came from our observation that inhibition of CCR1 signaling could fully restore sensitivity to Dex-induced apoptosis of partial GC-resistant cells (Fig. 4d). In this context, previous reports have linked CCR1 expression to decreased sensitivity to other anti-MM drugs^17,18^ and coupled pro-survival effects to CCR1 signaling^38^. Therefore, higher levels of CCR1 expression might well be associated with a more general drug-resistant phenotype.

We have tested the Dex-BX combination therapy in a myeloma xenograft mouse model to investigate the potential clinical relevance of our combination treatment beyond primary myeloma cells. We showed enhanced anti-MM effects of the combination compared to both monotherapies. However, due to poor solubility and pharmacokinetics of BX471 in mice^39^, high doses of compound and solvent (propylene glycol) had to be administered. Propylene glycol can cause nephrotoxicity when given repeatedly in mice^40^, which can explain the overall health concerns, weight loss and abnormalities observed during sacrifice in all treatment groups (see Supplementary Table 3 for a full report), since all animals received equal doses of propylene glycol. This limited the duration of the treatment to two days, which translated into rather moderate anti-MM effects observed *in vivo* compared to the substantial combination effects *in cellulo*. Further optimization with newer generations of CCR1 inhibitors could be a step forward, especially since CCX9588 and CCX721 have proven anti-myeloma effects in monotherapy^19,20^. However, despite the vast amount of CCR1 antagonists that have entered clinical trials for rheumatoid arthritis and multiple sclerosis, and were deemed safe and well-tolerated, none have entered the market due to a limited efficacy for said diseases^21^. So far, no CCR1 antagonist has been evaluated in a clinical trial for MM, despite its therapeutic or bone-sparing potential. This further emphasizes the need for alternative ways to target CCR1. Other than that, the redundancy of chemokines and their receptors provides another challenge^41^, even more as their expression patterns change during disease progression^42^ and during the development of therapy resistance^14^. Regarding the safety of the drug combination, further research is warranted as CCR1 is expressed on other hematopoietic cells^14^. Nonetheless, in a study by Ferguson et al., CCR1 was unbiasedly uncovered as an attractive new target for antigen-specific immunotherapy in myeloma, due to its distinctive expression profile on MM cells^14^.

Long term GC-use can lead to Dex-induced osteoporosis, resulting from decreased bone formation and increased bone marrow adiposity^43^. With osteolytic bone disease being a hallmark of MM and an ongoing therapeutic challenge, long-term GC-use in MM patients thus further compromises the already fragile bone. Currently bisphosphonates remain the golden standard for treatment of bone lesions, but care has to be taken to avoid renal toxicity^44^. The use of CCR1 inhibitors as an alternative treatment of myeloma bone disease, has previously been researched^13,19,37^, however, no studies have investigated the combination with other anti-MM drugs *in vivo*. Here, we show that beyond its anti- myeloma effects, Dex-BX combination also positively mitigated myeloma bone disease, as evidenced by higher serum OPG levels, which are inversely correlated with myeloma bone disease^29^. Future research should investigate the effect of Dex-BX combination treatment in a more advanced myeloma mice model that incorporates the complexity of the human bone as well^45^ or in a GC-sensitive immunocompetent mice model that displays osteolytic bone disease^46^. Such models allow studying the impact on Dex-induced osteoporosis, as our current mouse model and treatment duration did not allow to study this. These models also provide a better reflaection of the bone marrow microenvironment, which impacts the drug sensitivity of myeloma cells^10,11,15^.

We have characterized the proteome of MM1.S cells treated with Dex-BX to further understand the mechanisms of apoptosis induction by the drug combination. Pathway enrichment revealed a decrease in expression of lysosomal enzymes. As myeloma cells are plasma cells with a high protein turnover, they rely on autophagy to maintain homeostasis^47^. Autophagy can act as a double-edged sword in MM, either by inducing drug resistance and promoting tumor growth^48^, or by contributing to autophagic cell death^49^. We observed induction of autophagy by Dex as evidenced by lower levels of p62, in agreement with previous reports in MM cells^30^. However, whether this induction of autophagy is pro- cell death^30^ or is a way of MM cells to cope with the metabolic stress induced by Dex is unclear^50^. MM1.S cells treated with Dex-BX also had lower levels of p62, which indicates ongoing autophagy, yet, additionally the levels of CTSB (and other lysosomal proteases) are significantly lower in Dex-BX treated cells, while CTSB is essential for autophagic flaux^51^. A possible explanation is that, given the dependence of MM cells on autophagy, intervening in the clearance of the autophagic cargo by lysosomal proteases, might be a factor explaining the toxicity of the drug combination. The observation that chloroquine, an agent blocking lysosomal degradation, enhances the cytotoxicity of Dex, supports this hypothesis. Moreover, combinations of chloroquine with other anti-myeloma drugs showed potential *in vitro*^52^ and the combination of chloroquine with carfilzomib and Dex proved to be safe and tolerable in a Phase I clinical trial^53^. Besides the effect on lysosomal proteins, Dex-BX also shifts the balance between pro- (Bim) and antiapoptotic proteins (Mcl-1). Mcl-1 plays a pivotal role in survival of MM cells^33,34^ and was identified as a therapeutic vulnerability by an RNAi screening^54^. As such, selective Mcl-1 inhibitors have been developed such as S68345 of which we could confirm its cytotoxic effects in MM1.S and OPM-2 cells and of which a synergistic cell killing effect together with Dex was demonstrated before^55^. However, due to concerns regarding cardiotoxicity, clinical trials with these class Mcl-1 inhibitors were halted^56^. Dex-BX treatment could thus serve as an alternative route to lowering Mcl-1 levels in MM cells, without the toxicity concerns. Bim on the other hand, is crucial for GC-induced apoptosis of lymphoid cells^2^ and is further upregulated by Dex-BX in our experiments. Taken together, our data show a reduction in antiapoptotic Mcl-1 levels with Dex-BX combination, combined with an increase in Bim levels, thereby tipping the balance between pro- and antiapoptotic proteins into the direction of apoptosis.

Finally, a survival analysis based on expression of Dex-BX downregulated proteins, revealed that patients having low expression of the associated genes, had significantly better OS and PFS than patients with high expression. This finding gives a hint that the Dex-BX downregulated proteins are important for MM cell survival in a broader patient population and not only in the cell lines tested in this study. On the other hand, a likewise analysis done with proteins upregulated by Dex-BX, revealed that having high expression of these upregulated genes, was detrimental for patient survival. This indicates that some of the Dex-BX regulated proteins contribute to the cytotoxic effects of the combination, while others still contribute to MM cell survival. Feature selection could be applied to the dataset as a strategy to disentangle genes independently associated with survival from others^8^.

To conclude, our work complements previous studies done on targeting CCR1 for myeloma treatment, which mainly focused on mono-targeting CCR1. However, there is a growing body of evidence that combination of CCR1 antagonists with other anti-MM drugs holds promise. By investigating the combination of targeting CCR1 with GCs, still a cornerstone treatment of MM, we provide the basis for further pre-clinical and clinical research. We here found support for combination effects of both drugs *in cellulo, in vivo* and in primary patient material, resolved the action mechanisms of the drug combination, uncovered the potential association of CCR1 with GC resistance and proposed a GC resensitization strategy. With improvements of CCR1 antagonists, further preclinical validation can prelude clinical validation of CCR1 targeting combination therapies.

## Acknowledgements

This research was funded by Faculty of Medicine and Health Science of Ghent University. (AAP funding to BL). This work was further supported through VIB and Ghent University institutional funding to KDB and Ghent University institutional funding to SG. The authors thank Prof. Dr. Sven Eyckerman for his critical suggestions and support in generating the CCR1 overexpression plasmid. VIB Proteomics core for LC-MS/MS analysis and VIB Flow core for guidance in flaow cytometry experiments.

## Author Contributions

B.L.. D.C.. M.V.T.. P.V.V.. S.G.. K.D.B.. were involved in conceptualization and provided critical suggestions; B.L.. D.C. performed investigation and analysis; D.C.. F.D.. M.V.T.. E.S.. A.V.. S.T.S. provided investigational support; F.O. provided patient samples; S.E. provided critical suggestions for expression plasmid design; B.L.. D.C.. S.G.. K.D.B. wrote and reviewed the manuscript. All authors approved the final version of the manuscript.

## Competing interests

The authors declare no competing interests

## Data availability statement

LC-MS/MS data from Dex+BX471 treated MM1.S cells has been deposited to the ProteomeXchange Consortium via the PRIDE repository with the dataset identifier PXD055337. Publicly available datasets used in this study are the MMRF CoMMpass dataset (release IA14). which can be accessed through the MMRF Researcher Gateway (https://research.themmrf.org). Expression data from myeloma cell lines were obtained from The Keats lab (https://www.keatslab.org/data-repository). All other data described in this study are available within the source data file. data collection methods are described in the methods section.

## Notes

https://www.ebi.ac.uk/pride/login

### Competing Interest Statement

The authors have declared no competing interest.

