## Supplementary figures and methods for "CCR1 inhibition sensitizes multiple myeloma cells to glucocorticoid therapy"

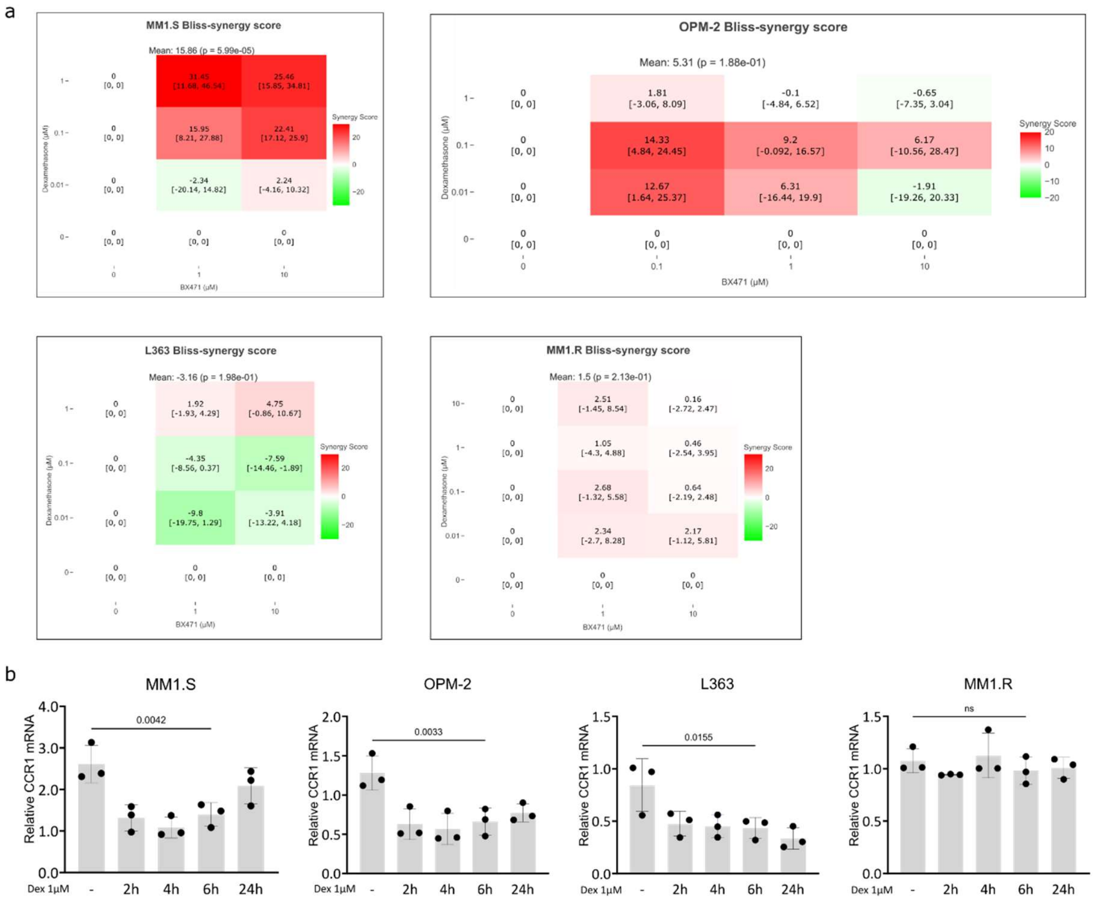


**Supplementary Figure 1** Bliss synergy scores for all drug concentration combinations tested in Fig. 1d and CCR1 mRNA levels in response to Dex.

**a** Bliss synergy scores for calculated for the different drug combinations shown in Fig. 1d. Mean Bliss synergy averaged over all concentration combinations is depicted above. A mean score >10 indicates a likelihood for drug synergy, a mean score <10 indicates a likelihood for additive drug effects. Reported p-values are for a t-test under the null hypothesis of drug independence. **b** Relative CCR1 mRNA levels after induction with 1 µM Dex for indicated timepoints, compared to solvent control (24h solvent treatment). 1-way ANOVA with post-hoc testing was performed and indicated are the *p*-values for the comparison between solvent control and 6h Dex 1 µM treatment.

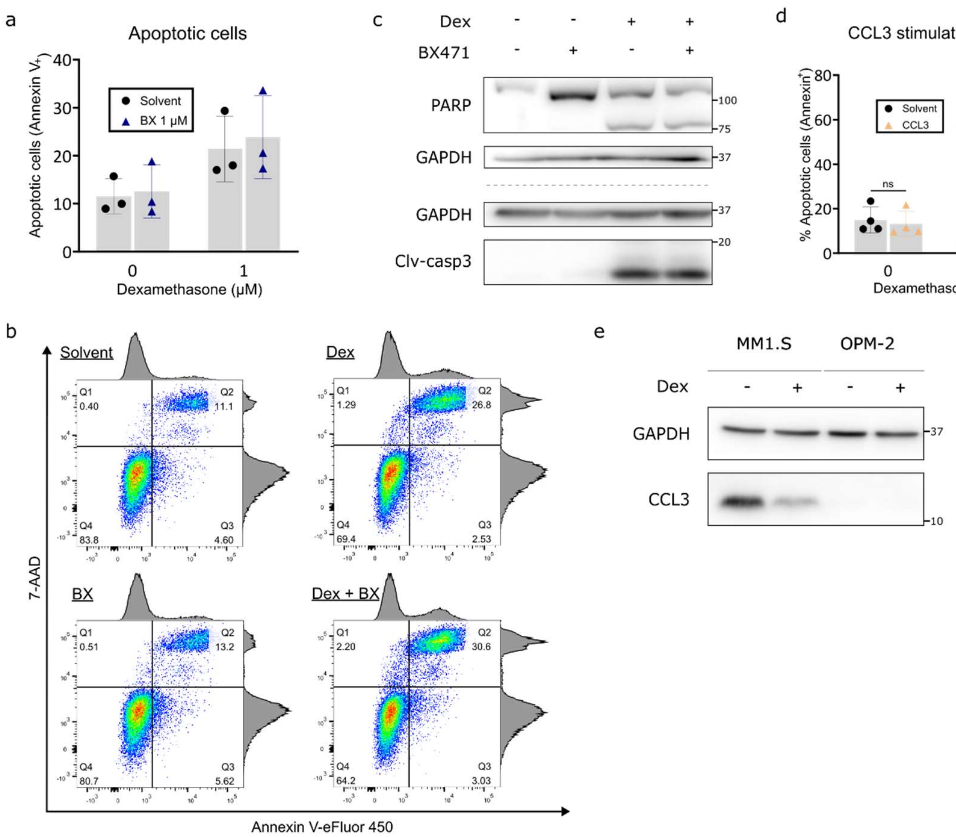


**Supplementary Figure 2** Dex-BX471 combination does not significantly increase Dex-induced apoptosis of OPM-2 cells, while supplementation of CCL3 tends to decrease sensitivity of OPM-2 cells to Dex-induced apoptosis.

**a** Apoptotic OPM-2 cells determined by flow cytometry after 24h treatment with 1 µM Dex (+ BX471 1 µM) (n=3). **b** Representative Annexin V/7-AAD plot after 24h Dex 1 µM (+ BX471 1 µM) treatment. **c** Western blot for apoptotic markers of OPM-2 whole cell lysates treated for 24h with 1 µM Dex (+ BX471 1 µM), GAPDH as loading control (n=3). **d** Apoptotic OPM-2 cells determined by flow cytometry after 48h treatment with Dex (1µM), in medium supplemented with 50 ng/mL recombinant CCL3 protein. The experiment was performed at the 48h timepoint instead of 72h with MM1.S cells, due to higher Dex-sensitivity of OPM-2 cells (Fig. 1e) **e** Western blot for CCL3 of MM1.S and OPM-2 whole cell lysates, treated for 24h with 1 µM Dex. GAPDH as loading control (n=3). Blot for MM1.S is the same as used in Fig. 2i. For **a** (n=3) and **d** (n=4) a 2-way ANOVA with post-hoc testing was performed with n.s. indicating the difference is not statistically significant.

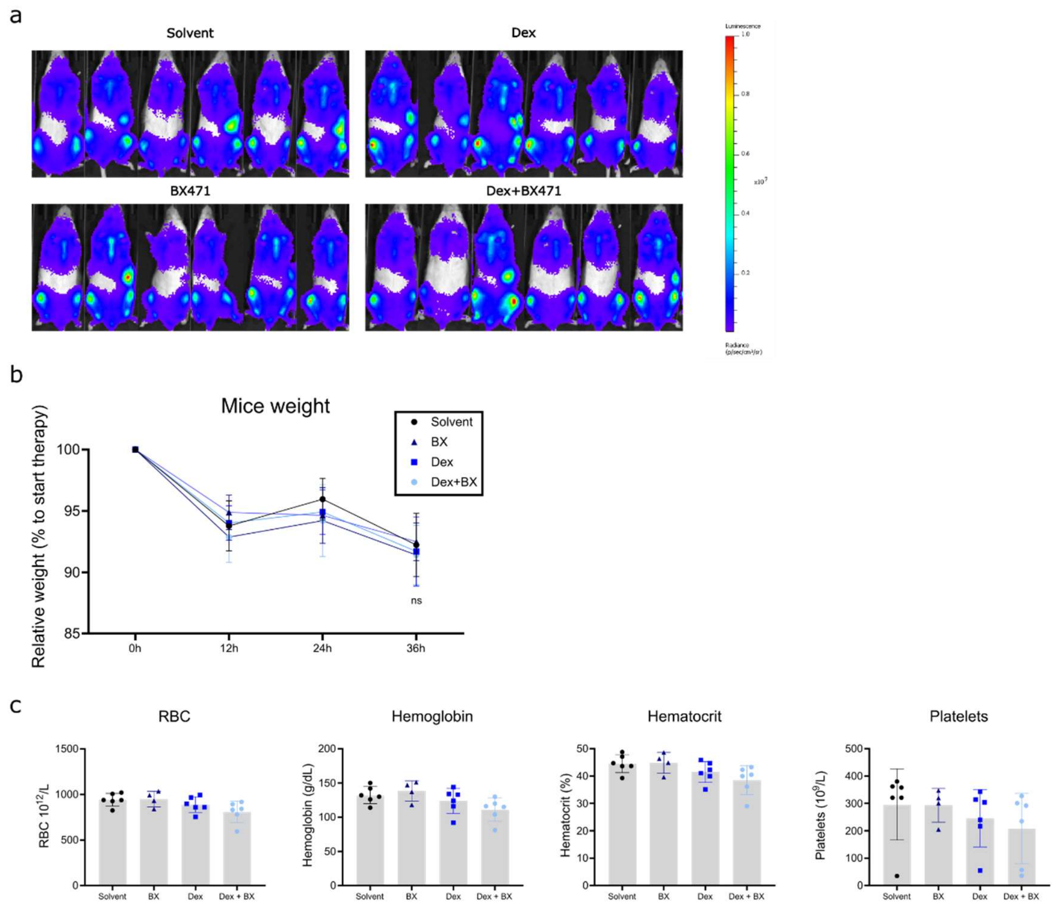


**Supplementary Figure 3** Tumor load before start of treatment, weight throughout treatment and blood parameters of mice at sacrifice.

**a** BLI images of mice 5-weeks after engraftment of MM1.S-luci cells at the start of treatment. BLI signal per mouse was quantified as average radiance per mouse (p/s/cm²/sr) and was used to randomize the mice in different treatment groups. **b** Weight of mice during the experiment, relative to start weight (n=6 mice per group). 2-way ANOVA with post-hoc testing was performed with no statistically significant differences between groups observed, indicated by ns at timepoint 36h. **c** Mouse blood parameters upon sacrifice, as measured by Vetscan HM5 Hematology analyzer. RBC = Red Blood Cell, Hct = hematocrit. N=6 mice per group, except for BX-group (n = 4), where 2 mice died during anesthesia of the last BLI where no whole blood could be collected. 1-way ANOVA with post-hoc testing was performed with no statistically significant differences between groups observed.

N=1

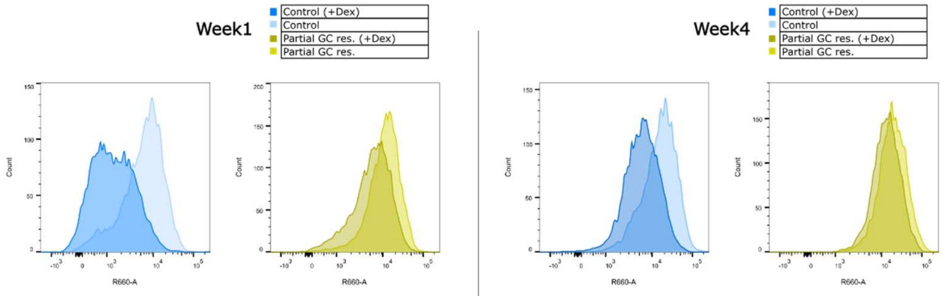


| Repetition | Week | Population | Dex 1 µM | Count | MFI CCR1-AF647 (R660-A) |
|---|---|---|---|---|---|
| N=1 | 1 | Control | - | 8424 | 9648 |
| N=1 | 1 | Control | + | 8515 | 2419 |
| N=1 | 1 | Partial GC res. | - | 8540 | 14859 |
| N=1 | 1 | Partial GC res. | + | 8510 | 8488 |

| Repetition | Week | Population | Dex 1 µM | Count | MFI CCR1-AF647 (R660-A) |
|---|---|---|---|---|---|
| N=1 | 4 | Control | - | 7800 | 22127 |
| N=1 | 4 | Control | + | 7740 | 8717 |
| N=1 | 4 | Partial GC res. | - | 8138 | 24804 |
| N=1 | 4 | Partial GC res. | + | 7917 | 17316 |

N=2

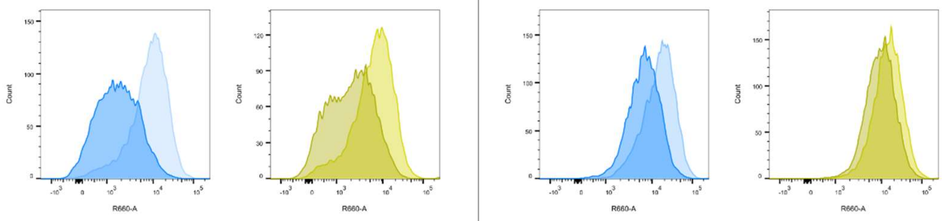


| Repetition | Week | Population | Dex 1 µM | Count | MFI CCR1-AF647 (R660-A) |
|---|---|---|---|---|---|
| N=2 | 1 | Control | - | 8269 | 12051 |
| N=2 | 1 | Control | + | 8198 | 2903 |
| N=2 | 1 | Partial GC res. | - | 8405 | 9494 |
| N=2 | 1 | Partial GC res. | + | 8359 | 3710 |

| Repetition | Week | Population | Dex 1 µM | Count | MFI CCR1-AF647 (R660-A) |
|---|---|---|---|---|---|
| N=2 | 4 | Control | - | 7927 | 20638 |
| N=2 | 4 | Control | + | 7703 | 9196 |
| N=2 | 4 | Partial GC res. | - | 8206 | 19129 |
| N=2 | 4 | Partial GC res. | + | 8115 | 12721 |

N=3

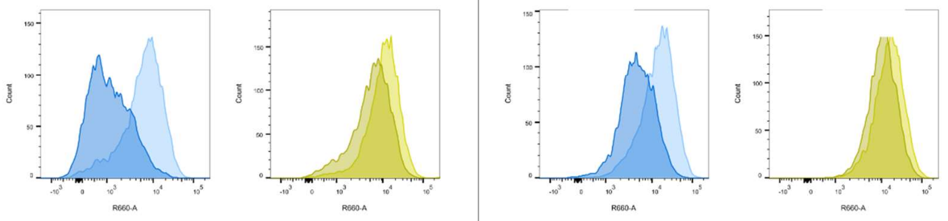


| Repetition | Week | Population | Dex 1 µM | Count | MFI CCR1-AF647 (R660-A) |
|---|---|---|---|---|---|
| N=3 | 1 | Control | - | 8489 | 9092 |
| N=3 | 1 | Control | + | 8758 | 2025 |
| N=3 | 1 | Partial GC res. | - | 8372 | 14405 |
| N=3 | 1 | Partial GC res. | + | 8475 | 8416 |

| Repetition | Week | Population | Dex 1 µM | Count | MFI CCR1-AF647 (R660-A) |
|---|---|---|---|---|---|
| N=3 | 4 | Control | - | 7671 | 19501 |
| N=3 | 4 | Control | + | 7166 | 6895 |
| N=3 | 4 | Partial GC res. | - | 7904 | 20308 |
| N=3 | 4 | Partial GC res. | + | 7859 | 13957 |

**Supplementary Figure 4** CCR1 plasma membrane expression levels in response to acute Dex-treatment throughout development of partial GC-resistance.

Flow cytometry plots of 3 independent biological replicates of the GC-resistance model at W1 and W4, showing CCR1 surface protein levels in response to a 24h treatment with 1 µM Dex. Cells were stained for cell surface CCR1 expression with a primary anti-CCR1 antibody and an AlexaFluor-647 labeled secondary antibody. Indicated is the mean fluorescence intensity (MFI) used to generate Fig. 4f. At W1 and at W4, the MFI of the control cells treated 24h with solvent control of the respective biological replicate served as the reference level for CCR1 expression.

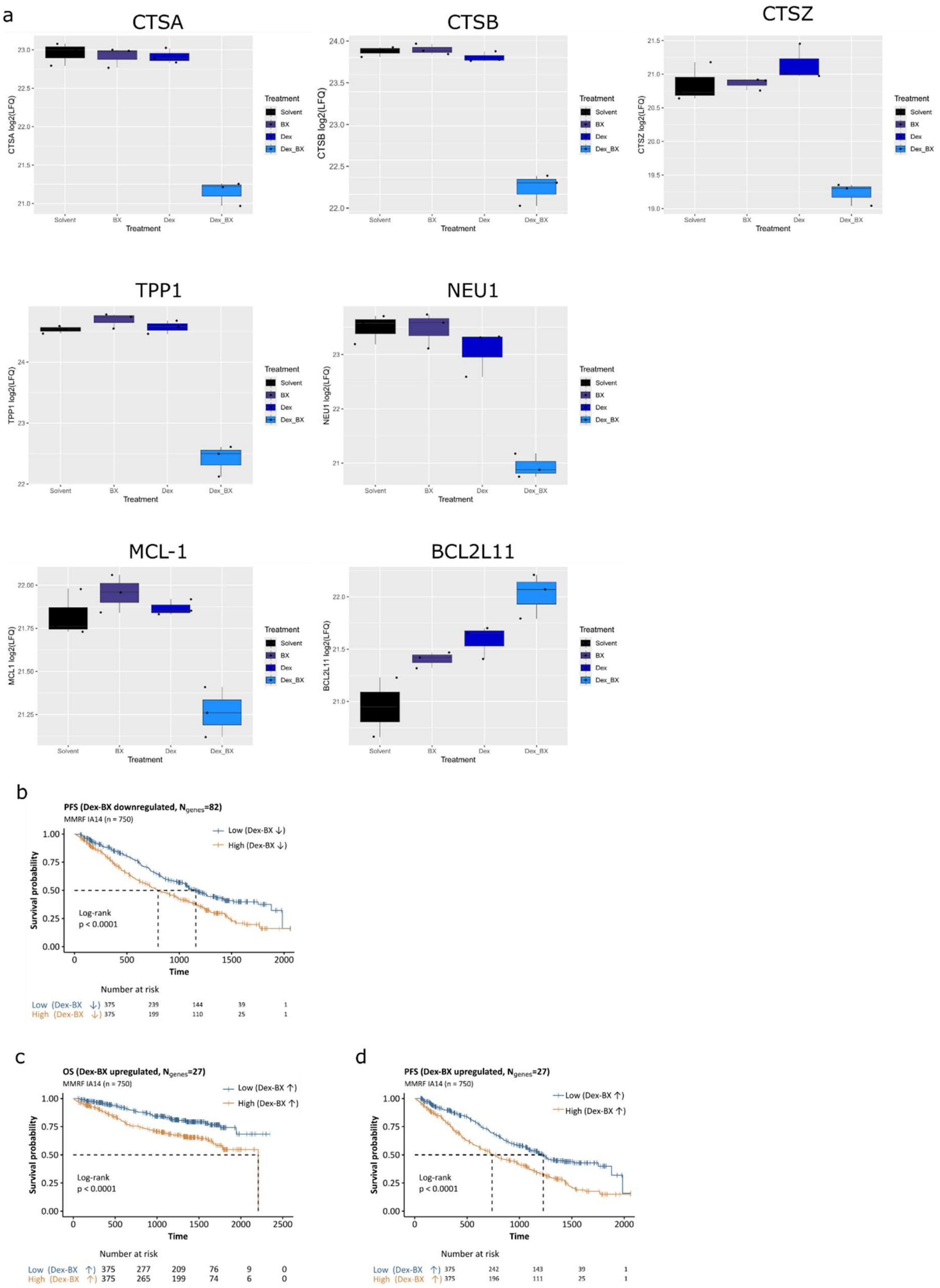


**Supplementary Figure 5** Raw LFQ-values for differentially expressed proteins by Dex:BX combination treatment, PFS and OS plots for Dex:BX combination treatment differentially expressed proteins

**a** LFQ-values of a selection of differentially expressed proteins by Dex:BX interaction. **b** PFS of patients included in the MMRF CoMMpass trial (IA14,n=750), stratified on expression of Dex:BX interaction downregulated genes in bone marrow samples of patients at diagnosis. **c** OS and **d** PFS of patients included in the MMRF CoMMpass trial (IA14, n = 750), stratified on expression of Dex:BX interaction upregulated genes in bone marrow samples of patients at diagnosis. Statistical significance was tested by a log-rank test with a null hypothesis of no difference in survival between both groups.

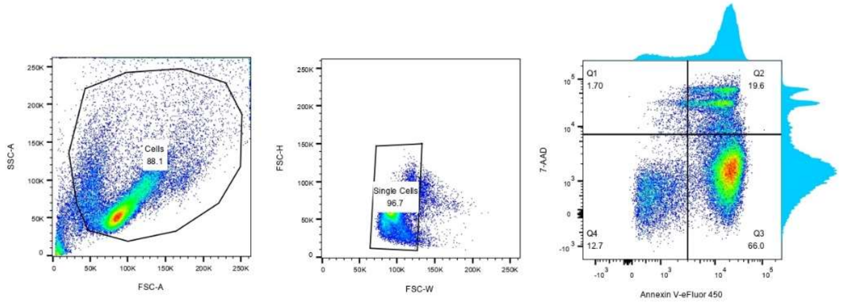


**Supplementary Figure 6** Flow cytometry gating strategy

**Supplementary Table 1**

Characteristics of patients included in the MMRF CoMMpass Trial (release IA14) who had RNA-sequencing of a bone marrow sample at diagnosis and/or at follow-up.

| Diagnosis (n=750) | |
|---|---|
| Characteristic | Number |
| Age (mean) | 64.03 (±10.84) |
| Male | 446 (59.5%) |
| Female | 304 (40.5%) |
| First line Dexamethasone | 708 (94.4%) |
| First line Prednisone | 32 (4.3%) |
| Other first line treatment | 10 (1.3%) |

| Progression (n=48) | |
|---|---|
| Characteristic | Number |
| Age (mean) | 67.15 (±10.36) |
| Male | 28 (58.3%) |
| Female | 20 (41.7%) |
| First line Dexamethasone | 40 (83.3%) |
| First line Prednisone | 7 (14.6%) |
| Other first line treatment | 1 (2.1%) |

**Supplementary Table 2**

Patient information for primary patient samples used in Fig. 1f. CTG = CelltiterGlo

| Pseudonym | Assay type, Treatment time | Gender | Age | Stage | M protein type |
|---|---|---|---|---|---|
| MM1 | CTG, 24h | M | 53 | MGUS | IgM kappa |
| MM2 | CTG, 48h | F | 31 | High risk SMM | IgG lambda |
| MM3 | CTG, 24h | F | 60 | newly diagnosed, prior to first therapy | IgG kappa |
| MM4 | CTG, 72h | F | 69 | newly diagnosed, prior to first therapy | IgG kappa |
| MM5 | CTG, 24h | F | 71 | Relapsed, prior to 2nd line of therapy | kappa light chain |
| MM6 | CTG, 72h | M | 56 | Relapsed, prior to 2nd line of therapy | IgA kappa |

### Supplementary Table 3

Mice health observations upon sacrifice.

| Cage | Mouse code | Group | Spleen weight | Mouse weight | Relative spleen weight | Remarks | Remarks 2 |
|---|---|---|---|---|---|---|---|
| 1 | 1_B | Solvent | 0.06 | 23 | 0.0026 | | |
| 1 | 1_2L | Solvent | 0.156 | 22.3 | 0.0070 | | |
| 2 | 2_2L | Solvent | 0.069 | 23.1 | 0.0030 | Liver not bright red | Liquid/air in intestines |
| 4 | 4_2L | Solvent | 0.154 | 19 | 0.0081 | Liver not bright red | Liquid/air in intestines (more than others) |
| 5 | 5_1L | Solvent | 0.103 | 20.8 | 0.0050 | | |
| 5 | 5_2L | Solvent | 0.176 | 20 | 0.0088 | Liquid/air in intestines | |
| 1 | 1_2R | Dex | 0.063 | 21.3 | 0.0030 | | |
| 3 | 3_B | Dex | 0.111 | 22.3 | 0.0050 | Liquid/air in intestines | |
| 3 | 3_1L | Dex | 0.133 | 20.6 | 0.0065 | Liver not bright red | |
| 3 | 3_1R | Dex | 0.055 | 21.4 | 0.0026 | Liver not bright red | |
| 4 | 4_B | Dex | 0.034 | 22.9 | 0.0015 | Liver not bright red | |
| 4 | 4_1L | Dex | 0.044 | 21.3 | 0.0021 | | |
| 1 | 1_1L | BX | 0.041 | 23 | 0.0018 | Liquid/air? in intestines | Died during last BLI anaesthesia |
| 1 | 1_1R | BX | 0.14 | 22 | 0.0064 | Liquid/air? In intestines | Died during last BLI anaesthesia |
| 2 | 2_B | BX | 0.039 | 22.7 | 0.0017 | Liver not bright red + spot on liver | Liquid/air in intestines |
| 2 | 2_2R | BX | 0.128 | 19.5 | 0.0066 | Liquid/air in intestines | |
| 3 | 3_2L | BX | 0.16 | 18.5 | 0.0086 | Liver not bright red | Liquid/air in intestines (less than others) |
| 5 | 5_B | BX | 0.117 | 20 | 0.0059 | Liver not bright red | Liquid/air in intestines (more than others) |
| 2 | 2_1L | Dex_BX | 0.035 | 24.1 | 0.0015 | | |
| 2 | 2_1R | Dex_BX | 0.021 | 24.4 | 0.0009 | (Digested) blood in intestines and stomach | Bleeding in stomach |
| 3 | 3_2R | Dex_BX | 0.05 | 21.8 | 0.0023 | Liver not bright red | Liquid/air in intestines |
| 4 | 4_1R | Dex_BX | 0.038 | 23.7 | 0.0016 | Liver not bright red | |
| 4 | 4_2R | Dex_BX | 0.024 | 23 | 0.0010 | | |
| 5 | 5_1R | Dex_BX | 0.07 | 18.2 | 0.0038 | Liver not bright red | |

**Supplementary Table 4**

Differentially expressed proteins by Dex-BX combination treatment compared to solvent (model without interaction term)

See separate file: Supplementary table 4_Differentially expressed proteins_Dex-BX_No interaction.xlsx

**Supplementary Table 5**

Differentially expressed proteins by Dex:BX combination treatment (model with interaction term)

See separate file: Supplementary table 5_Differentially expressed proteins_DexBX interaction term.xlsx

**Supplementary Table 6**

KEGG 2021 enriched terms in the Dex:BX differentially expressed proteins list (Supplementary Table 5)

| Term | Overlap | P.value | Adjusted.P.value | Odds.Ratio | Combined.Score | Genes |
|---|---|---|---|---|---|---|
| Other glycan degradation | 12/202 | 4.16E-04 | 0.0323 | 13.846 | 107.781 | MANBA;GLB1;NEU1;ENGASE |
| Pyrimidine metabolism | 10/142 | 9.53E-04 | 0.0338 | 5.833 | 40.575 | NT5E;DUT;RRM1;RRM2;TK1; TYMS |
| Lysosome | 12/128 | 1.25E-05 | 0.0029 | 5.088 | 57.456 | CTSA;MANBA;GLB1;NEU1;CTSZ; PPT1;AP4S1;TPP1;AP3S1; GLA;LGMN;CTSB |
| Cell cycle | 11/124 | 4.78E-05 | 0.0056 | 4.776 | 47.516 | RB1;CCNA2;CCNB2;CCNB1;WEE1; ORC6;SMAD3;CHEK1;PLK1; BUB1;MAD2L1 |
| Progesterone-mediated oocyte maturation | 8/100 | 1.02E-03 | 0.0338 | 4.239 | 29.214 | CCNA2;CCNB2;CCNB1;PLK1;KIF22; BUB1;AURKA;MAD2L1 |
| Apoptosis | 10/142 | 6.81E-04 | 0.0338 | 3.704 | 27.011 | DFFA;PARP4;TRADD;CTSZ;HTRA2; BIRC5;CFLAR;SPTAN1;MCL1;CTSB |
| Epstein-Barr virus infection | 12/202 | 9.43E-04 | 0.0338 | 3.094 | 21.557 | RB1;CCNA2;CD40;MAVS;CR2; PSMC6;PSMC1;TRADD;VIM;ITGAL; JAK3;NFKBIB |

### Supplementary Table 7

Primary and secondary antibodies used in WB and flow cytometry

| Primary antibodies | | |
|---|---|---|
| **Target** | **Ref.** | **Supplier** |
| Bim | 2933 | Cell Signaling Technology |
| CCL3 | ab259372 | Abcam |
| CCR1 | MAB145-SP | R&D Systems |
| Cleaved-caspase 3 | 9664 | Cell Signaling Technology |
| CTSB | 31718 | Cell Signaling Technology |
| GAPDH | G8795 | Sigma |
| GR | sc-393232 | Santa Cruz Biotechnology |
| LC3 | L8918 | Sigma |
| Mcl-1 | 5453 | Cell Signaling Technology |
| PARP | 556494 | BD Biosciences |
| p62 (SQSTM1) | P0067 | Sigma |

| Secondary antibodies | | |
|---|---|---|
| **Target** | **Ref.** | **Supplier** |
| anti Mouse IgG HRP | NA931 | GE Healthcare |
| anti Rabbit IgG HRP | NA934 | GE Healthcare |
| anti Mouse-Alexa Fluor 647 | A-21235 | Invitrogen |

### Supplementary Table 8

Primers and shRNA sequences

| Primer | | |
|---|---|---|
| **Target** | **FW sequence** | **REV sequence** |
| CCR1 | AGGCAAAAGGAAGCAGGGTT | AAGTCCAAGATGGCAGTCGG |
| RPL13A | CCTGGAGGAGAAGAGGAAAGAGA | TTGAGGACCTCTGTGTATTTGTCAA |
| SDHA | TGGGAACAAGAGGGCATCTG | CCACCACTGCATCAAATTCATG |
| YWHAZ | ACTTTTGGTACATTGTGGCTTCAA | CCGCCAGGACAAACCAGTAT |

| shRNA | |
|---|---|
| **Target** | **Sequence** |
| Control | CCGGCAACAAGATGAAGAGCACCAACTCGAGTTGGTGCTCTTCATCTTGTTGTTTTT |
| shCCR1 _1 | CCGGCCCTACAATTTGACTATACTTCTCGAGAAGTATAGTCAAATTGTAGGGTTTTT |
| shCCR1 _2 | CCGGGCTCTGAAACTGAACCTCTTTCTCGAGAAAGAGGTTCAGTTTCAGAGCTTTTT |

### Supplementary methods:

#### Patient survival analysis

Data from the MM research foundation (MMRF) CoMMpass trial (release IA14) was used, accessed via the MMRF research portal (https://research.themmrf.org). RNA-sequencing data as normalized TPM gene expression values together with clinical data served as input data for survival analysis, which was performed using 'Survival' package in R (version 3.5-8). Statistical significance was calculated by a log-rank test. Expression data was filtered to only obtain RNA-sequencing data from BM samples (n = 750). Overall survival (OS) was defined as the time from diagnosis until death from any cause or until the latest time point the patient was known to be alive, in which case the patient was censored. For survival analysis based on gene expression levels at diagnosis, patients were divided into 2 equally sized groups, based on their average z-score normalized expression data ranked from low to high. For 49/750 patients, a subsequent sample (=progression sample) was taken for RNA-sequencing, after having received at least one line of therapy. Only patients with detectable CCR1 expression both at diagnosis and at progression, were included for further analysis (48 patients). For survival analysis on gene expression after at least one line of therapy, patients were divided into 2 groups based on CCR1 gene expression levels at progression, relative to CCR1 expression at diagnosis. The cutoff was made at patients having ($CCR1_{TPM}$ – levels at progression) / ($CCR1_{TPM}$ – levels at diagnosis) ≥ 2 (25/48 patients).

#### Bliss Synergy scores

Synergy scores were calculated as described by Ianevski et al.(1), through the web application http://synergyfinder.fimm.fi . The Bliss synergy score can be interpreted as excess drug response above expectation, due to drug interactions. Reported p-values are for a t-test under the null hypothesis of drug independence. Scores >10 indicate a drug interaction that is likely synergistic (Synergyfinder documentation available at https://synergyfinder.fimm.fi/synergy/synfin_docs/#datanal).

#### Shotgun proteomics and data analysis

MM1.S cells were treated for 24h (n=3 biological replicates), subsequently washed and cell pellets were stored at -80°C. Mass spectrometry sample preparation and data analysis were performed as described below. The resulting protein levels were expressed as log2(LFQ) values. Subsequent data analysis was done in R using the limma package (version 3.58.1). A linear model with either 3 factors (Dex, BX and Dex+BX) or a model with 2 factors (Dex and BX) an interaction term (Dex:BX) was fitted to the data. Statistical testing for differential expression is done by moderated t-tests, with p-values adjusted for multiple testing ($p_{adj}$) by the method of Benjamini and Hochberg(2,3). The mass spectrometry proteomics data have been deposited to the ProteomeXchange Consortium via the PRIDE partner repository with the dataset identifier PXD055337 (accessible via https://www.ebi.ac.uk/pride/)

*Sample preparation*

Cell pellets were homogenized in 100 µl lysis buffer containing 5% sodium dodecyl sulphate (SDS) and 50 mM triethylammonium bicarbonate (TEAB). pH 8.5. Next. the resulting lysate was transferred to a 96-well PIXUL plate and sonicated with a PIXUL Multisample sonicator (Active Motif) for 5 minutes with default settings (Pulse 50 cycles. PRF 1 kHz. Burst Rate 20 Hz). After centrifugation of the samples for 15 minutes at 2.204xg at room temperature (RT) to remove insoluble components. the protein concentration was measured by bicinchoninic acid (BCA) assay (Thermo Scientific) and from each sample 100 µg of protein was isolated to continue the protocol. Proteins were reduced and alkylated by addition of 10 mM Tris(2-carboxyethyl)phosphine hydrochloride and 40 mM chloroacetamide and incubation for 10 minutes at 95°C in the dark. Phosphoric acid was added to a final concentration of 1.2% and subsequently samples were diluted 7-fold with binding buffer containing 90% methanol in 100 mM TEAB. pH 7.55. The samples were loaded on the 96-well S-Trap plate (Protifi). placed on top of a deepwell plate. and centrifuged for 2 min at 1.500 x g at RT. After protein binding. the S-trap plate was washed three times by adding 200 µl binding buffer and centrifugation for 2 min at 1.500 x g at RT. A new deepwell receiver plate was placed below the 96-well S-Trap plate and 50 mM TEAB containing 1 µg trypsin (1/100. w/w) was added for digestion overnight at 37°C. Using centrifugation for 2 min at 1.500 x g. peptides were eluted in three times. first with 80 µl 50 mM TEAB. then with 80 µl 0.2% formic acid (FA) in water and finally with 80 µl 0.2% FA in water/acetonitrile (ACN) (50/50. v/v). Eluted peptides were dried completely by vacuum centrifugation. Samples were dissolved in 100 µl 0.1% TFA and desalted on reversed phase (RP) C18 OMIX tips (Agilent). The tips were first washed 3 times with 100 µl pre-wash buffer (0.1% TFA in water/ ACN (20:80. v/v)) and pre-equilibrated 5 times with 100 µl of wash buffer (0.1% TFA in water) before the sample was loaded on the tip. After peptide binding. the tip was washed 3 times with 100 µl of wash buffer and peptides were eluted twice with 100 µl elution buffer (0.1% TFA in water/ACN (40:60. v/v)). The combined elutions were transferred to HPLC inserts and dried in a vacuum concentrator.

*LC-MS/MS analysis*

Peptides were re-dissolved in 20 µl weak wash solvent (WW) (0.1% trifluoroacetic acid in water/acetonitrile (ACN) (99.5:0.5. v/v)) of which 2µl was injected for LC-MS/MS analysis on an Vanquish™ Neo UHPLC System in-line connected to a Orbitrap Exploris 240 mass spectrometer (Thermo). Injection was performed in trap-and-Elute workflow in combined Control mode (maximum flow of 60 µl/min and maximum pressure of 800 bar) in WW on a 5 mm trapping column (Thermo scientific. 300 µm internal diameter (I.D.). 5 µm beads). The peptides were separated on a 250 mm Aurora Ultimate. 1.7µm C18. 75 µm inner diameter (Ionopticks) kept at a constant temperature of 45°C. Peptides were eluted by a gradient starting at 0.5 % MS strong wash solvent (SW) (0.1% FA in acetonitrile) reaching 26% MS SW in 75 min. 44% MS SW in 95 min. 56% MS SW in 100 minutes followed by 5-minute wash at 56% MS SW and column equilibration in Pressure Control mode (separation column: fast equilibration. maximum pressure of 1500 bar. equilibration factor=2; Trap column: fast wash and equilibration. wash factor=100) with MS WW . The flow rate was set to 300 nl/min.

The mass spectrometer was operated in data-independent mode. automatically switching between MS and MS/MS acquisition. Full-scan MS spectra ranging from 400-900 m/z with a normalized target value of 300%. a maximum fill time of 25 ms and a resolution at of 60.000 were followed by 30 quadrupole isolations with a precursor isolation width of 10 m/z for HCD fragmentation at an NCE of 30% after filling the trap at a normalized target value of 2000% for maximum injection time of 45 ms. MS2 spectra were acquired at a resolution of 15.000 with a scan range of 200-1800 m/z in the Orbitrap analyser without multiplexing. The isolation intervals were set from 400 – 900 m/z with a width of 10 m/z using window placement optimization.

EASY-IC™ was used in the start of the run as Internal Mass Calibration and QCloud has been used to control instrument longitudinal performance during the project.

*Data analysis*

Analysis of the mass spectrometry data was performed in DiaNN (version 1.8.1). Precursor false discovery rate was set at 1%. Spectra were searched against the Homo sapiens protein sequences in the Uniprot database (database release version of January 2024). containing 20.597 sequences (www.uniprot.org). Enzyme specificity was set as C-terminal to arginine and lysine. also allowing cleavage at proline bonds with a maximum of 2 missed cleavages. Variable modifications were set to oxidation of methionine residues and acetylation of protein N-termini while fixed modification was set to carbamidomethylation of cystein residues. Matching between runs was enabled. Mainly default settings were used. except for the addition of a 400-900 m/z precursor mass range filter and MS1 and MS2 mass tolerance was set to 10 and 20 ppm respectively.

Further data analysis of the results was performed with an in-house script in the R programming language. Protein expression matrices were prepared as follows: the DIA-NN main report output table was filtered at a precursor and protein library q-value cut-off of 1 % and only proteins identified by at least one proteotypic peptide were retained. After pivoting into a wide format. iBAQ intensity columns were then added to the matrix using the DIAgui's R package get_IBAQ function. LFQ intensities were log2 transformed and replicate samples were grouped. Proteins with less than 3 valid values in at least one group were removed and missing values were imputed from a normal distribution centered around the detection limit (package DEP) leading to a list of 7.479 quantified proteins in the experiment. used for further data analysis. Considering Dex+BX as a different treatment. protein abundance between pairs of sample groups was compared (BX vs Solvent; Dex vs Solvent; Dex_BX vs Solvent;) and statistical testing for differences between two group means was performed. using the package limma. Statistical significance for differential regulation was set to a false discovery rate (FDR) of $< 0.05$ and $|log2FC| = 1$. Considering Dex:BX as an interaction term between Dex and BX treatments. protein abundance between treatments (Solvent. Dex. BX and Dex:BX) was compared and statistical testing for differences between groups was performed. using the package limma. Statistical significance for differential regulation was set to a false discovery rate (FDR) of $< 0.05$ and $|log2FC| = 1$. Results are presented in Supplementary Tables 4 and 5. Z-scored LFQ intensities from significantly regulated proteins were plotted in a heatmap after non-supervised hierarchical clustering.
